# Structure-based mapping of the TβRI and TβRII receptor binding sites of the parasitic TGF-β mimic, Hp-TGM

**DOI:** 10.1101/2020.12.08.416701

**Authors:** Ananya Mukundan, Chang-Hyeock Byeon, Cynthia S. Hinck, Danielle J. Smyth, Rick M. Maizels, Andrew P. Hinck

**Author notes:** Corresponding Author: Prof. Andrew P. Hinck, Department of Structural Biology, University of Pittsburgh School of Medicine, Biomedical Science Tower 3, Room 1035, 3501 Fifth Avenue, Pittsburgh, PA 15260, U.S.A.

## Abstract

TGF-β is a secreted signaling protein involved in many physiological processes: organ development, production and maintenance of the extracellular matrix, as well as regulation of the adaptive immune system. As a cytokine, TGF-β stimulates the differentiation of CD4^+^ T-cells into regulatory T-cells (T_regs_) that act to promote peripheral immune tolerance. The murine parasite *Heligmosomoides polygyrus* takes advantage of this pathway to induce inducing Foxp3^+^ T_regs_ in a similar manner using a TGF-β mimic (TGM), comprised of five tandem complement control protein (CCP) domains, designated D1-D5. Despite having no structural homology to TGF-β or to TGF-β family proteins, TGM binds directly to the TGF-β type I and type II receptors, TβRI and TβRII. To further investigate, NMR titration, and SPR and ITC binding experiments were performed, showing that TGM-D2, with the aid of D1, binds TβRI and TGM-D3 binds TβRII. Competition ITC experiments showed that TGM-D3 competes with TGF-β for binding to TβRII, consistent with TGM-D3-induced NMR chemical shift perturbations of TβRII which aligned with the solvent inaccessible areas of TβRII upon binding TGF-β. Thus, TGM-D3 binds to the same edged β-strand of TβRII that is used to bind TGF-β. Competition ITC experiments demonstrated that TGM-D1D2 and TGF-β3:TβRII compete for binding to TβRI, while TGM-D2-induced NMR chemical shift perturbation of TβRI showed that TGM-D2 binds to the same pre-helix extension of TβRI as does the TGF-β/TβRII binary complex. The solution structure of TGM-D3 revealed that while it has the overall structure of a CCP domain, TGM-D3 has an insertion in the hypervariable loop uncommon to CCP domains. These findings suggest that parasitic TGM, despite its lack of structural similarity to TGF-β, evolved to take advantage of the binding regions of the mammalian TGF-β type I and type II receptors. The structure of this TGM domain, along with the predicted structure of other *H. polygyrus* secreted proteins reported in the literature, suggest that TGM is part of a larger family of evolutionarily-adapted immunomodulatory CCP-containing proteins.

## Introduction

Helminth parasites are extraordinarily prevalent in all host organisms and remain major human health burdens in tropical regions of the world^1,2^; their longevity reflects an evolutionarily-refined ability to evade the immune system through multiple molecular strategies which are only now becoming understood^3-5^. A number of helminth infections are associated with activation of the regulatory T-cell (T_reg_) compartment which dampens inflammation and restricts anti-parasite immunity, either through expansion of the host’s pre-existing T_regs_ or through inducing *de novo* differentiation of peripheral T-cells into the T_reg_ subset^6^. Notably, infection with the mouse model helminth *Heligmosomoides polygyrus* expands T_reg_ activity, and worm clearance can be induced by antibody-mediated depletion of T_regs_^7^. T_regs_ express a defining transcription factor, Foxp3, which can be induced in naive T-cells by the pleiotropic cytokine TGF-β^8-10^. Moreover, *H. polygyrus* secretory-excretory products (HES) can induce T_regs_ by signaling through the TGF-β receptors, TβRI and TβRII^11^. Fractionation and analysis by a TGF-β response bioassay isolated the protein in HES responsible for stimulating the TGF-β signaling pathway and inducing T_regs_, as a five-domain ca. 420 amino acid protein designated TGF-β mimic (TGM), which is fully active as a full-length protein requiring no post-translational processing, and bearing no sequence similarity to any member of the TGF-β family^12^.

Canonical TGF-β proteins control a multitude of pathways in cellular differentiation^13-15^ and immune homeostasis^9,13,16^, and are tightly regulated by pro-domain processing to yield ∼110-amino acid cystine-knotted monomers tethered together into active 25kDa homodimers by a single inter-chain disulfide bond. They signal by assembling a heterotetrameric complex with two pairs of two serine/threonine kinase type I and type II receptors, TβRI and TβRII respectively^17-19^. TGF-β-dependent differentiation of naïve CD4^+^ cells into CD4^+^ CD25^+^ Foxp3^+^ T_regs_, is essential for peripheral immune tolerance^8,9^ as mice lacking TGF-β1 exhibit perinatal mortality and develop multi-organ inflammatory disease and die after maternal TGF-β is depleted upon weaning^13^. Dysregulation of the TGF-β signaling pathway has been implicated in the pathogenesis of several human diseases, ranging from inflammatory bowel disease^20^, renal and cardiac fibrosis^21,22^, and many soft tissue cancers^21,23,24^. In the lattermost setting, TGF-β dysregulation prevents effective checkpoint immunotherapy^25,26^ and thus offers a therapeutic target in its own right^27^.

In contrast to the single-domain structure of mature TGF-β, the parasitic-encoded TGM molecule is composed of 5 modular domain, all with distinct similarity to the complement control protein (CCP) famliy; CCP domains are ca. 60 amino acids in length and are comprised of multiple short β-strands tethered together by two highly conserved disulfide bonds (Cys^I^-Cys^III^, Cys^II^-Cys^IV^)^28^. CCP domains are not usually found alone, but are instead are usually found as arrays, in some cases with as many as 30 repeated modules^28^. These domains are present in numerous proteins, but they are most prevalent in the family of proteins that act to regulate complement (RCA), which among others include decay accelerating factor (DAF), Factor H (FH), and Complement C3b/C4b Receptor 1 (CR1)^28^. In *H. polygyrus*, more than 30 CCP-containing proteins have been detected^29,30^, including HpARI (*H. polygyrus* Alarmin Release Inhibitor) and HpBARI (*H. polygyrus* Binds Alaarmin Receptor and Inhibits) which suppress IL-33 signaling by binding IL-33 and its receptor ST-2^31-33^; these cytokines prime innate and adaptiive type 2 immue responses and thus suppression via HpARI and HpBARI acts as an alternatae form of immunosuppression. Similar to TGM, Hp-ARI and Hp-BARI contain multiple CCP domainsn (3 and 2, respectively) and have significant insertions not present in canonical CCP domains^12,31,32^.

Previous studies have shown that of the five domains of TGM, only the first three, D1-D3, are required for TGF-β signaling activity^29^. Previous studies also showed that TGM, in contrast to TGF-β, binds TβRI with high affinity (K_D_ 52.1nM) and TβRII with lower affinity (K_D_ 0.55 μM) and that it does so without the pronounced inter-dependence of TβRI and TβRII binding characteristic of the TGF-βs^17,34^. Here we biophysically characterized the individual domains of TGM to show that binding of TGM to human TβRI and TβRII is modular in nature, with D1-D2 and D3 binding to TβRI and TβRII with 1:1 stoichiometry, respectfully. We characterized the binding sites that TβRI and TβRII use to bind TGM using ITC, SPR, and NMR finding that they utilize structural motifs similar to those used to bind TGF-β. Finally, we determined the solution structure of TGM-D3 which showed that TGM-D3 takes on the overall fold of a CCP domain with two key differences: 1) loops replacing two β-strands, and 2) an atypical insertion at the hypervariable loop (HVL). Using NMR chemical shift perturbations, the binding site of TβRII on TGM-D3 was mapped, with large targeted shifts across the four β-strands, but particularly focused on the C-terminal β-strands. The atypical insertion at the HVL dramatically increases its length, which allows it to block both faces of the N-terminal region of TGM-D3 from binding TβRII. Thus, TβRII is able to insert the edged β-strand it uses to bind TGF-β alongside the face of TGM-D3, interacting most with the C-terminal β-strands of TGM-D3. This highlights that the parasite has evolved to take advantage of host TGF-β receptors, and that more broadly *H. polygyrus* has adapted its own CCP domain-containing proteins for the purpose of protein mimicry and host immunomodulation.

## Results

### Isolation and folding of TGM-D1, -D2, and -D3

Previously, *in vitro* TGF-β bioassays demonstrated that only TGM domains 1-3 were required for full TGF-β Smad reporter activity and induction of CD4^+^ CD25^+^ Foxp3^+^ T_regs_^29^. Truncation of domains 4 and 5 (TGM-D1-3) showed minimal reduction in TGF-β activity, while loss of domain 1 (TGM-D2-5), domains 3-5 (TGM-D1-2), or domains 2-5 (TGM-D1) completely abolished TGF-β activity^29^. TGM was furthermore shown to require both TβRI and TβRII to elicit TGF-β signaling, as TGM activity was inhibited by Sβ431542^35^, a TβRI kinase inhibitor, and by ITD-1 which stimulates ubiquitin-dependent degradation of TβRII ^12^.

The engagement of the TGF-β receptors by TGM was thus investigated by assessing the receptor binding properties of TGM domains 1, 2, and 3, each as an isolated protein. To enable isotopic labeling, the individual domains were produced in *E. coli*. TGM-D1, -D2, and -D3 each contain four cysteines and are expected to form two disulfides by homology to CCP domains. Though steps were taken to produce each domain as a soluble protein by fusion to thioredoxin and expression at reduced temperature (18 °C), this was not successful. Thus, the fusion proteins were expressed at 37 °C in the form of insoluble inclusion bodies, and refolded in the presence of a glutathione redox couple.

The isolated proteins were validated by mass spectrometry, which showed that their masses matched, to within 0.5 Da or less of the mass calculated assuming the formation of two disulfide bonds (i.e. loss of two protons for each disulfide bond). To further validate the refolded proteins, they were labeled with ^15^N and analyzed by recording ^1^H-^15^N HSQC spectra in phosphate buffer at pH 6.0 at 37 °C. The CCP family of proteins, to which TGM belongs, are characterized by four or more β-strands^36,37^, and thus in addition to having the expected number of peaks indicative of a homogenous pairing of the cysteines, the labeled proteins would also be expected to have well-dispersed spectra, with minimal clustering in the random coil region (between 7.8 and 8.3 ppm in the ^1^H dimension). This was observed for TGM-D3, which had very close to the expected number of backbone amide resonances (80 observed, 81 expected) and excellent signal dispersion (Fig. S1A).

TGM-D2, although also exhibiting excellent signal dispersion, had more backbone amide resonances than expected (106 observed, 76 expected), suggesting sample heterogeneity (Fig. S1B). This could be due to heterogenous pairing of the disulfides, or may result from conformational dynamics. To distinguish between these, HSQC ZZ-exchange spectra were recorded with mixing times ranging between 0 – 250 ms ^38^. These experiments identified at least 12 pairs of peaks undergoing exchange on this timescale, indicating that the protein is likely homogenous with respect to the pairing of its cysteines, but undergoes a slow conformational transition that leads to two conformations in solution. The process responsible for the doubling was not investigated, but would be consistent with proline *cis*:*trans* isomerization owing to the slow (ca. 100 ms) timescale by which the exchange cross peaks build up and the fact that TGM-D2 has four additional proline residues relative to TGM-D1 (Fig. S1C, Table S1).

TGM-D1, in contrast to TGM-D2 and TGM-D3, had poor signal dispersion, with most peaks clustered in the random coil region of the spectrum (Fig. S2A). This narrow dispersion might reflect the presence of non-native protein due to mispairing of cysteines, or it might be due to other reasons, such as aggregation. To investigate the latter, increasing concentration of CHAPS was added to the NMR buffer and the protein concentration was decreased. This led to the appearance of a large number of peaks outside of the random coil region (Fig. S2B-D). The spectrum with 20 μM TGF-D1 and 10 mM CHAPS in the buffer had roughly the expected number of peaks (85), but as well a few intense peaks in the random coil region of the spectrum (Fig. S2D). Thus, TGM-D1 appears to be natively folded, but is not entirely disaggregated under these conditions.

### NMR-detected binding of TGM-D1, -D2, and -D3 by TβRI and TβRII

To assess binding of TβRII by TGM-D1, TGM-D2, and TGM-D3, unlabeled TβRII was titrated into samples of ^15^N-labeled TGM-D1, TGM-D2, or TGM-D3. The addition of TβRII resulted in significant perturbations in more than half of the backbone amide signals of TGM-D3 (Fig. 1A), but little to no perturbations in the signals of either TGM-D1 or TGM-D2 (Fig. S3A-B). Through the course of the titration of TGM-D3 with TβRII, peaks corresponding to both the bound and unbound forms of TGM-D3 were observed at intermediate titration points (1:0.35 and 1:0.70), indicative of slow-exchange binding (Fig. 1C). This suggested that TGM-D3 binds to TβRII with relatively high affinity, but weakly or not at all to TGM-D1 or TGM-D2. The presence of 10 mM CHAPS in the TGM-D1 sample may, however, impede binding and thus a role of TGM-D1 in binding TβRII cannot at this point be excluded. The NMR titrations of TGM-D3 into TβRII further suggest that TGM-D3 binds TβRII with a stoichiometry of 1:1 (Fig. 1C).

**Figure 1.**
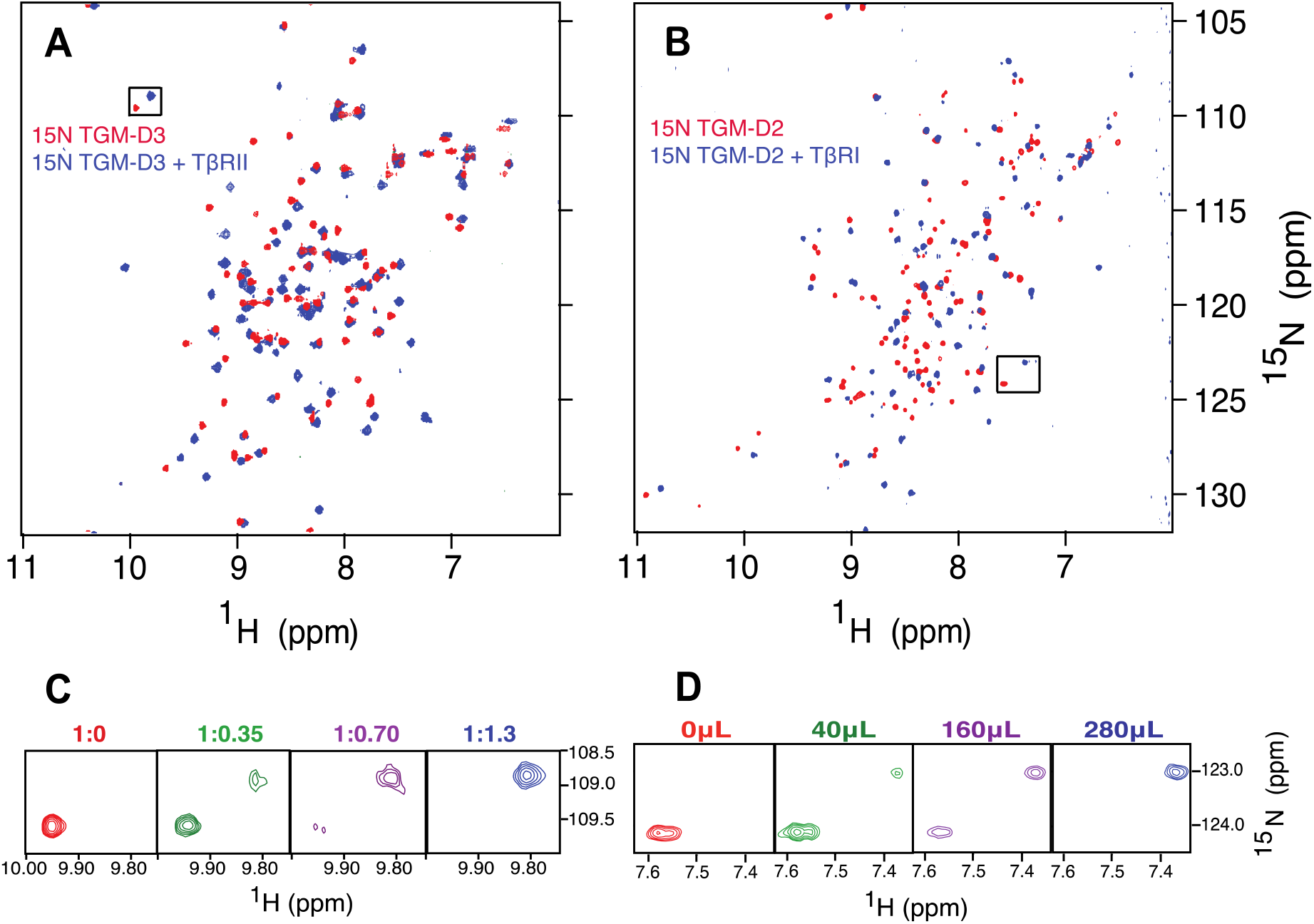
Binding of TGM-D2 and TGM-D3 by TβRI and TβRII, respectively. A-B. ^1^H-^15^N HSQC spectra of 0.02 mM ^15^N TGM-D3 (A) or 0.2 mM ^15^N TGM-D2 (B) alone (red) overlaid with the spectrum of the same sample, but with an excess of unlabeled TβRII (A) or TβRI (B) added (blue). All spectra were recorded in 25 mM sodium phosphate, 50 mM sodium chloride, 5% ^2^H_2_O pH 6.0 (A,C) or pH 7.0 (B,D), 310 K. Expansion of the boxed region of the spectra in panels A and B with all titration points is shown below in panels C and D, respectively, where numbers indicate either molar equivalents of TβRII added (C) or volume of TβRI stock added (μL) (D).

The addition of TβRI resulted in significant perturbations in more than half the backbone amide signals of TGM-D2 (Fig. 1B), but little to no perturbations in the signals of either TGM-D1 or TGM-D3 (Fig. S3C-D). However, similar to TGM-D1:TβRII, the presence of 10 mM CHAPS in the TGM-D1 sample may impede binding and thus a role of TGM-D1 in binding TβRI cannot be excluded. Through titrations in which increasing amounts of TβRI were added to TGM-D2, peaks corresponding to both the bound and unbound forms were observed under sub-stoichiometric conditions (Fig. 1D), indicative of relatively high affinity binding. The stoichiometry of binding was confirmed by titrating the amount of TβRI needed to fully convert the unbound form of ^15^N TGM-D2 to the bound form using NMR (Fig. S4A-D), with samples from each titration analyzed by size-exclusion chromatography (SEC) (Fig. S4E-I) Furthermore, a sample of TβRI with excess TGM-D2 was analyzed by SEC as monitored by multiangle light scattering (SEC-MALS) (Fig. S4J). The peak corresponding to the complex had a molecular mass of 17.1 ± 1.5 kDa, while the later eluting peak, corresponding to TGM-D2, had a mass of 10.1 ± 0.4 kDa. These experimentally measured masses are close to those expected for a 1:1 TβRI:TGM-D2 complex (18.8 kDa) or TGM-D2 alone (9.3 kDa), thus TGM-D2 binds TβRI with 1:1 stoichiometry. The titration of ^15^N TGM-D2 with TβRI leads to resolution of the conformational doubling in the spectrum of TGM-D2 that was previously noted (Fig. S5A-B). Thus, TGM-D2 appears to be primarily responsible for binding TβRI, and binding stabilizes TGM-D2 in one of its two native conformations.

To confirm primary binding of TGM-D2 to TβRI and binding of TGM-D3 to TβRII, and to assess the potential role for TGM-D1, the converse NMR titration experiments were performed, with samples of ^15^N-labeled TβRI and TβRII being titrated with unlabeled TGM-D1, -D2, or -D3, all in buffers lacking CHAPS. The titration of ^15^N TβRII with unlabeled TGM-D3 resulted in significant perturbations in its backbone amide signals (Fig. S6A), but titration with TGM-D1 or TGM-D2 did not (Fig. S7A,B). The titration of ^15^N TβRII with TGM-D3 further revealed the simultaneous appearance of peaks corresponding to the unbound and bound at sub-stoichiometric ratios, confirming that TGM-D3 is the main binding partner for TβRII and that it binds with high affinity (Fig S6C). The absence of any shifts upon titration of ^15^N TβRII with TGM-D1 suggests the lack of binding previously observed was not due to interference by CHAPS.

The titration of ^15^N TβRI with unlabeled TGM-D2 supports the prior results with a significant perturbation in over half of the backbone amide signals (Fig. S6B). There are peaks corresponding to both the unbound and bound forms at intermediate titration points, confirming that binding is high-affinity (Fig. S6D). The addition of unlabeled TGM-D3 into ^15^N TβRI resulted in little to no perturbation of the backbone amide signals, confirming that TGM-D3 does not bind TβRI (Fig. S8A). Titration of ^15^N TβRI with unlabeled TGM-D1 in the absence of CHAPS resulted in the weakening of many, and full disappearance, of nearly a third of the TβRI backbone signals, along with weak perturbations of other residues (Fig. S8B). The disappearance of some of the TβRI signals and small shifts of other residues is likely because it is binding to TGM-D1 and being incorporated into a TGM-D1 aggregate. Thus, although the nature of the binding precludes detailed analysis of the interaction, TGM-D1 does appear to bind TβRI in the absence of CHAPS and TGM-D1 may contribute to its binding by TGM.

### ITC and SPR quantification of T/JRII binding

To determine whether TGM-D3 is the only domain responsible for binding TβRII, or whether TGM-D1 or -D2 might also play a role, ITC binding experiments were performed in which either full-length TGM (TGM-FL) or individual domains of TGM, were titrated into TβRII in the calorimetry cell. The strong exothermic response for TGM-FL and TGM-D3 with TβRII indicate binding (Fig. 2C,D). The fitting of the integrated heat to a standard binding isotherm for both binding reactions yielded K_D_s and enthalpies of 0.55 μM (0.26 – 1.08, 68.3% CI))/ -7.14 kcal mol^-1^ (−7.80 – −6.55, 68.3% CI) and 1.15 μM (0.87 – 1.52, 68.3% CI)/ −10.65 kcal mol^-1^ (−11.22 – −10.14, 68.3% CI) for TGM-FL and TGM-D3, respectively (Fig. 2E-F, Table 1). Titration of TβRII with TGM-D1 or TGM-D2, in contrast, led to weak endothermic and exothermic responses, which were observed in buffer only titrations, indicating that neither TGM-D1 nor TGM-D2 binds TβRII (Fig. 2A-B, right inset). These observations, along with the similarity of the binding affinities of TGM-FL and TGM-D3 to TβRII, indicate that TGM-D3 is responsible for nearly all or all of the binding capacity of TGM-FL for TβRII.

**Figure 2.**
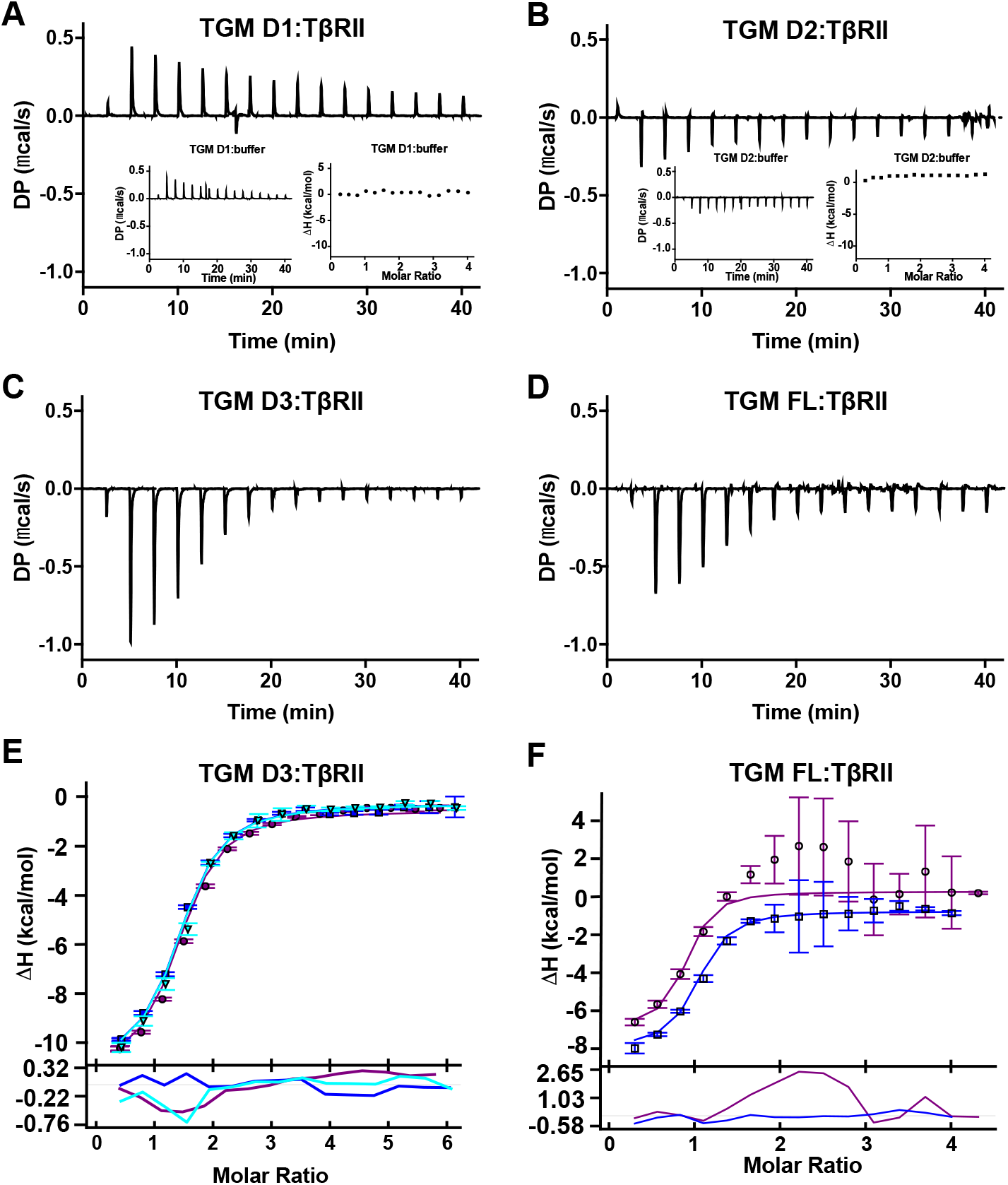
Binding of TβRII by TGM-D1, -D2, -D3, and -FL as assessed by ITC. A-D. Thermograms obtained upon injection of TGM-D1 (A), TGM-D2 (B), TGM-D3 (C), or TGM-FL (D) into TβRII. The insets shown on the left-hand side in panels A and B correspond to thermograms for injection of TGM-D1 or TGM-D2 into buffer (left). The insets shown on the right hand side in panels A and B correspond to subtraction of the integrated heats for the TGM-D1:buffer or TGM-D2:buffer titration from the TGM-D1:TβRII or TGM-D2:TβRII titration. E-F. Integrated heats for the titrations shown in panels C and D together with 1 and 2 replication titrations (E and F, respectively) with residuals as a function of the TGM domain relative to TβRII. The data points correspond to the integrated heats and the colored lines correspond to a global fit to a 1:1 binding model.

**Table 1.**
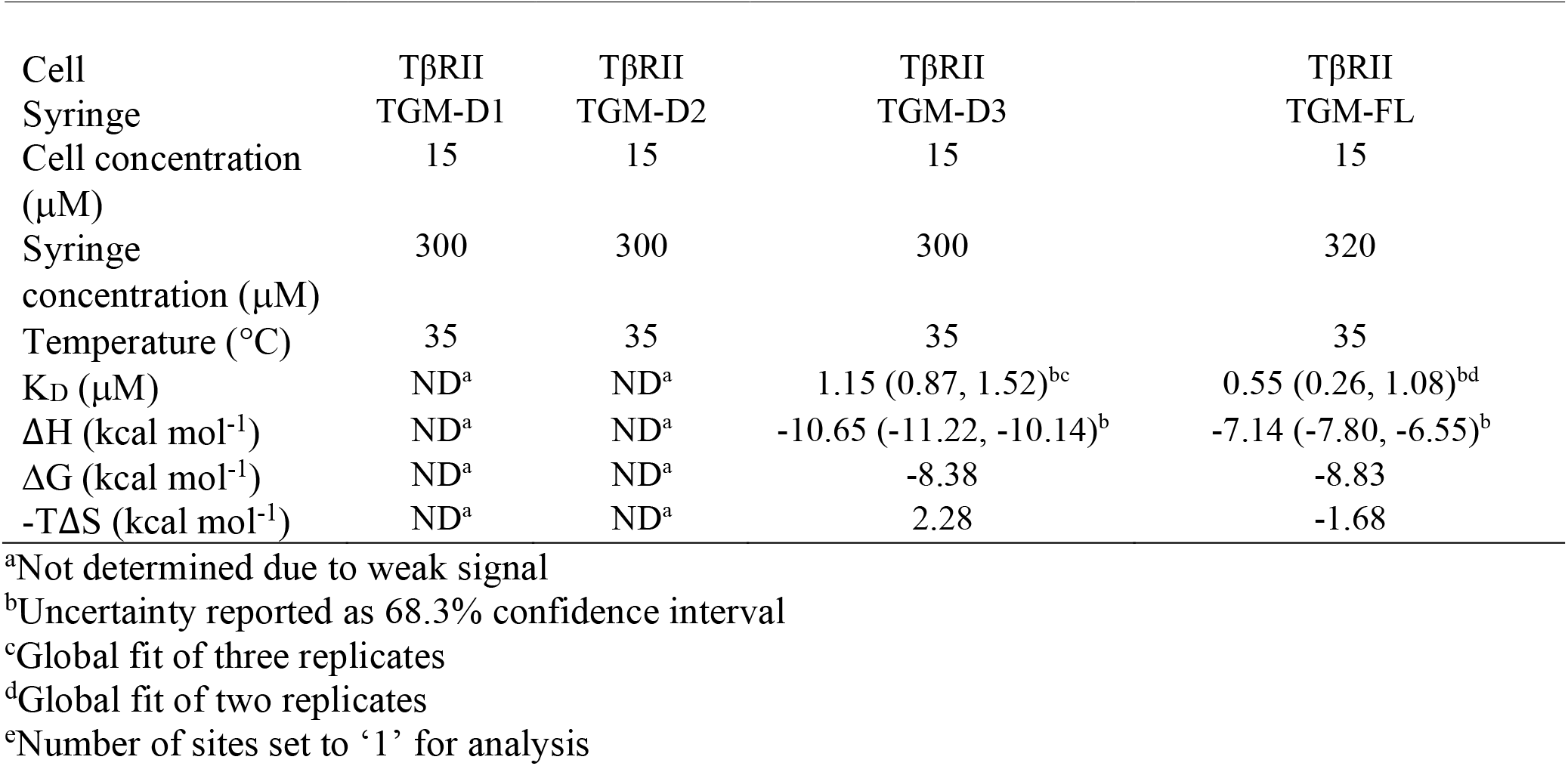
TGM:TPRII binding as assessed by ITC.

The results above were further validated by SPR measurements in which TGM-FL, or individual domains of TGM, were injected over biotinylated avitagged TβRII captured on a streptavidin-coated sensor chip. These injections yielded robust concentration-dependent responses when TGM-FL or TGM-D3 were injected over the immobilized TβRII, but not when TGM-D1, TGM-D2, or a construct that included both TGM domains 1 and 2, TGM-D1D2, were injected (Fig. 3). The K_D_ values derived by fitting the TGM-FL and TGM-D3 sensorgrams to a simple (1:1) kinetic model were comparable, 0.61 ± 0.01 μM and 0.91 ± 0.02 μM, respectively, though as with the ITC data, TGM-FL demonstrated slightly greater affinity than TGM-D3 for TβRII (Table S3, Fig. 3D-E). The SPR results are therefore consistent with the overall conclusion derived from both the NMR and ITC experiments, that TGM-D3 was responsible for all or almost all of the binding capacity of TGM-FL for TβRII.

**Figure 3.**
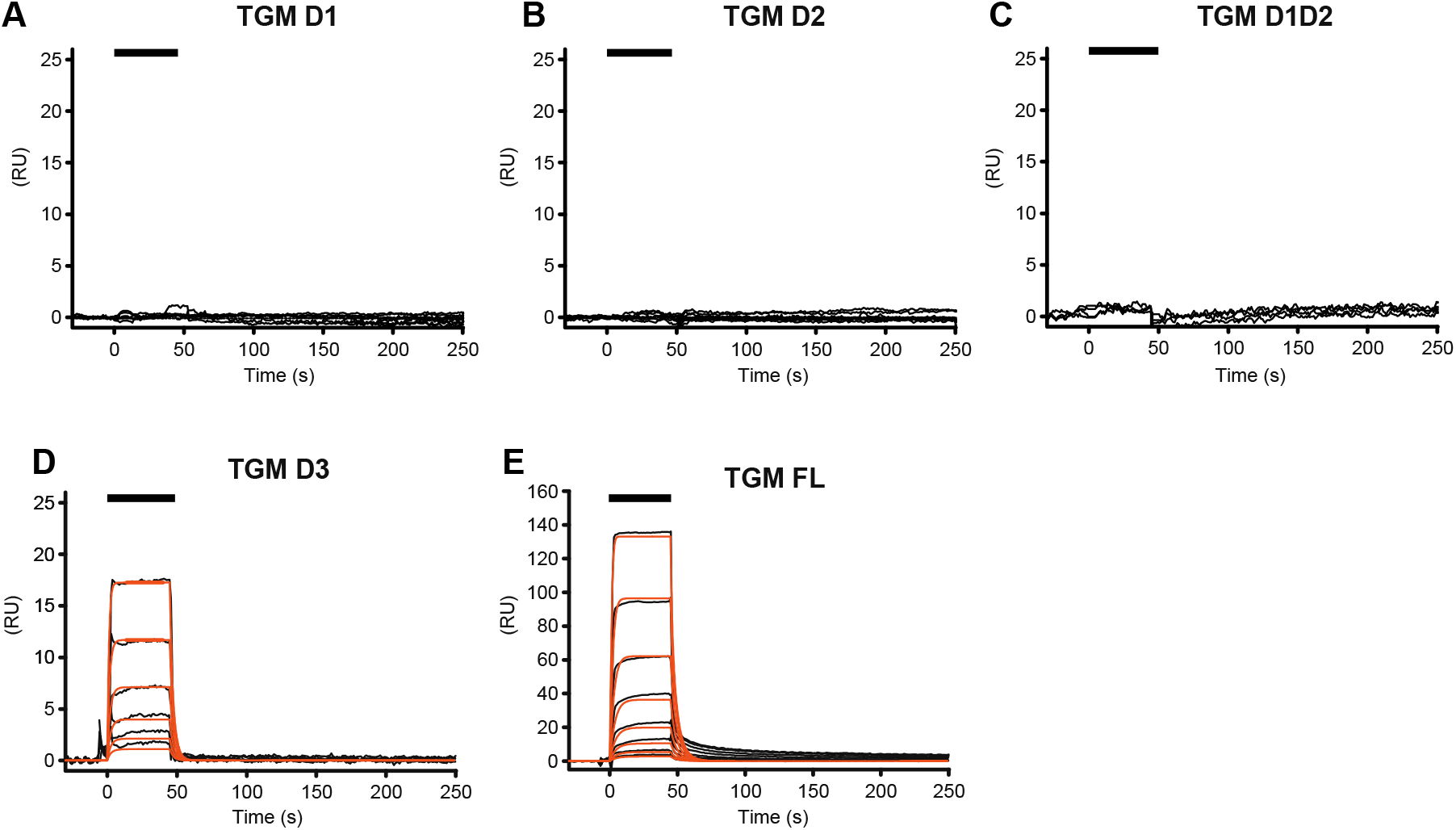
Binding of TPRII by TGM-D1, TGM-D2, TGM-D3, TGM-D1D2, and TGM-FL as assessed by SPR. A-E. SPR sensorgrams obtained upon injection of TGM-D1 (A), TGM-D2 (B), TGM-D1D2 (C), TGM-D3 (D), or TGM-FL (E) over immobilized TBRII. Sensorgrams, obtained upon injections of a two-fold dilution series of each TGM construct (TGM-D1 15.6 – 500 nM, TGM-D2 15.6 – 500 nM, TGM-D3 15.6 – 500 nM, TGM-D1D2 15.6 – 500 nM, and TGM-FL 15.6 – 500 nM) are shown in black, with the fitted curves in orange (data for TGM-D1, TGM-D2, and TGM-D1D2 were not fit due to weak signal). Black bars shown in the upper left specify the injection period.

### ITC and SPR quantification of T/JRI binding

Similarly, to determine whether TGM-D2 is the only domain responsible for binding TβRI, or whether TGM-D1 or -D3 might also play a role, ITC binding experiments were performed in which either TGM-FL, or individual domains of TGM were titrated into TβRI in the calorimetry cell. The strong exothermic response for TGM-FL and TGM-D2 with TβRI indicate binding (Fig. 4C, E, F,H). The fitting of the integrated heat to a standard binding isotherm for both binding reactions yielded K_D_s and enthalpies of 52.1nM (29.3 – 90.2, 68.3% CI)) and −16.69 kcal mol^-1^ (−18.26 – −15.35, 68.3% CI) and 1.47μM (0.45 – 4.58, 68.3% CI) and −17.66 kcal mol^-1^ (−26.95 – −13.18, 68.3% CI) for TGM-FL and TGM-D2, respectively (Fig. 4F,H) Table 2). Titration of TβRI with TGM-D1 or TGM-D3, in contrast, led to a weak endothermic (TGM-D1) or no response (TGM-D3) similar to that of an injection into buffer, and no response, indicative of no binding or weak binding. The finding that TGM-D2 bound with an affinity about thirty-fold weaker than TGM-FL complements the previous NMR titration data suggesting that TGM-D1 might also contribute to binding TβRI. To test this, a construct that included both TGM-D1 and TGM-D2 (TGM-D1D2), was produced in bacteria, refolded, and purified to homogeneity. This construct was titrated into TβRI and yielded a robust concentration-dependent response (Fig. 4D, G). The K_D_ and enthalpy was 25.3nM (10.7 – 48.3, 68.3% CI) and −18.70 kcal mol^-1^ (−19.59 – −17.85, 68.3% CI) respectively, which was comparable to that of TGM-FL. These observations indicate that both domains TGM-D1 and TGM-D2 are required to recapitulate the binding capacity of TGM-FL for TβRI.

**Figure 4.**
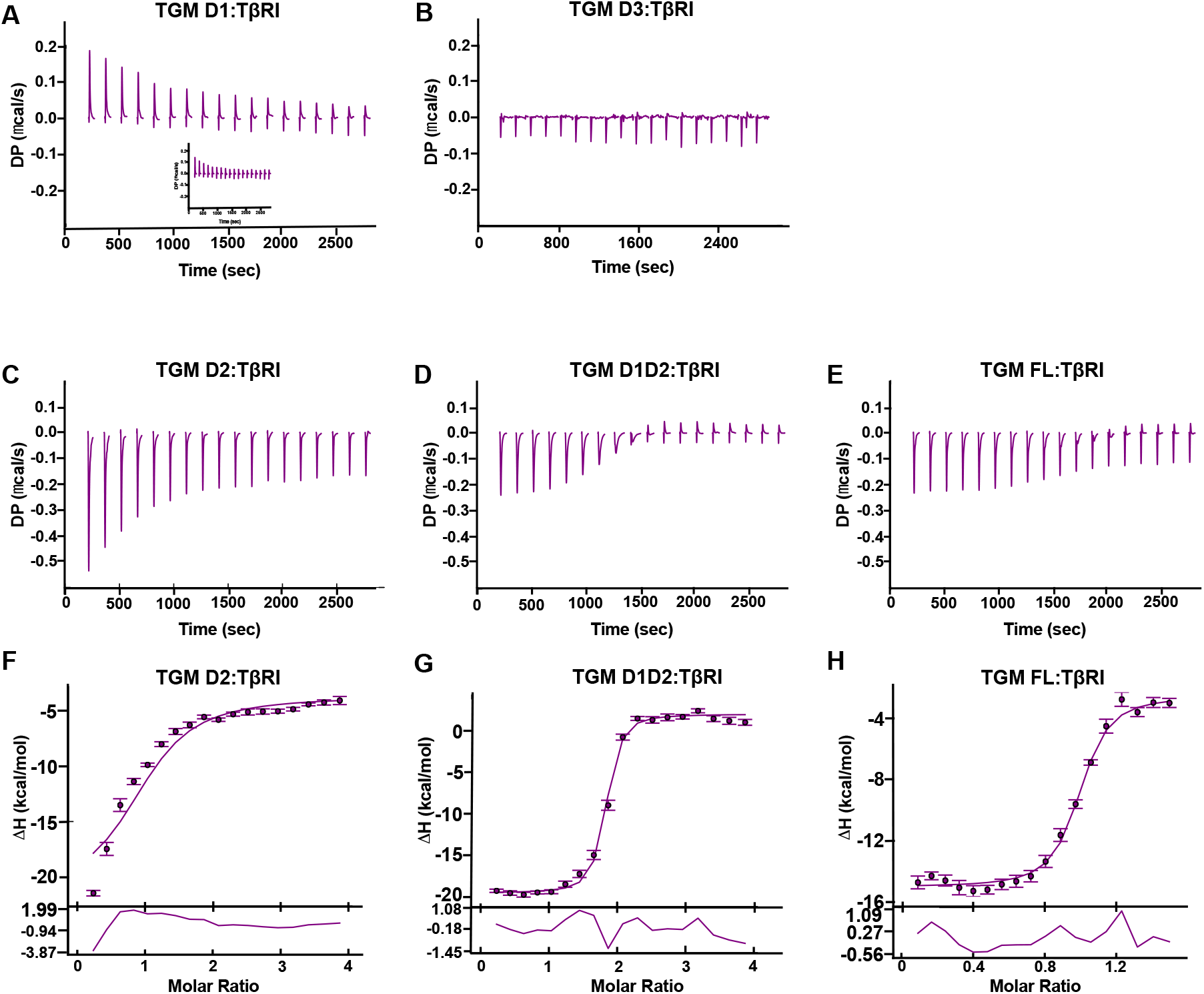
Binding of TβRI by TGM-D1, −D2, -D3, -D1D2, and -FL as assessed by ITC. A-E. Thermograms obtained upon injection of TGM-D1 (A), TGM-D3 (B), TGM-D2 (C), TGM-D1D2 (D) or TGM-FL (E) into TβRI. The insets shown in panel A corresponds to subtraction of the integrated heats for the TGM-D1:buffer titration from the TGM-D1:TBRI titration. F-H. Integrated heats for the titrations shown in panels C-E (F-H respectively) with residuals. The data points correspond to the integrated heats and the purple lines correspond to a fit to the model for 1:1 binding.

**Table 2.** TGM:TPRI binding as assessed by ITC.

| Cell | TβRI | TβRI | TβRI | TβRI | TβRI |
| --- | --- | --- | --- | --- | --- |
| Syringe | TGM-D1 | TGM-D2 | TGM-D3 | TGM-D1D2 | TGM-FL |
| Cell concentration (μM) | 7.5 | 7.5 | 7.5 | 7.5 | 7.5 |
| Syringe concentration (μM) | 150 | 150 | 135 | 100 | 58 |
| Temperature (°C) | 25 | 25 | 25 | 25 | 25 |
| K<sub>D</sub> (nM) | ND<sup>a</sup> | 1470 (451 – 4580)<sup>bc</sup> | ND<sup>a</sup> | 25.3 (10.7, 48.3)<sup>bc</sup> | 52.1 (29.3 – 90.2)<sup>bc</sup> |
| ΔH (kcal mol<sup>-1</sup>) | ND<sup>a</sup> | -17.66 (-26.95 – -13.18)<sup>b</sup> | ND<sup>a</sup> | -18.70 (-19.59 – -17.85)<sup>b</sup> | -16.69 (-18.26 – -15.35)<sup>b</sup> |
| ΔG (kcal mol<sup>-1</sup>) | ND<sup>a</sup> | -7.96 | ND<sup>a</sup> | -10.63 | -9.94 |
| -TΔS (kcal mol<sup>-1</sup>) | ND<sup>a</sup> | 9.70 | ND<sup>a</sup> | 8.34 | 6.75 |

<sup>a</sup>Not determined due to weak signal<sup>b</sup>Uncertainty reported as 68.3% confidence interval<sup>c</sup>Fit for one replicate<sup>d</sup>Number of sites set to '1' for analysis

SPR experiments were also performed with TβRI, TGM-FL, TGM-D1D2, or individual domains of TGM, injected over biotinylated avi-tagged TβRI captured on a streptavidin-coated sensor chip. These injections yielded robust concentration-dependent responses when TGM-FL, TGM-D2, or TGM-D1D2 were injected over the immobilized TβRI, but not when TGMs-D1 or TGM-D3 were injected (Fig. 5). The K_D_ values derived by fitting the TGM-FL and TGM-D1D2 sensorgrams to a simple (1:1) kinetic model were comparable, 13.1 ± 0.4 nM and 24.1 ± 0.1 μM, respectively, though TGM-FL demonstrated slightly greater affinity than TGM-D1D2 for TβRI (Table S4, Fig. 5D-E). The K_D_ derived from kinetic analysis of the TGM-D2 sensorgram was and 309 ± 4 nM (Fig. 5B, Table S4), indicating that the affinity of TGM-D2 for TβRI is approximately 25 times weaker than the TGM-FL, consistent with the previous ITC results. The SPR results are therefore consistent with the overall conclusion derived from both the NMR and ITC experiments that both TGM-D1 and -D2 are required for the full binding capacity of TGM for TβRI.

**Figure 5.**
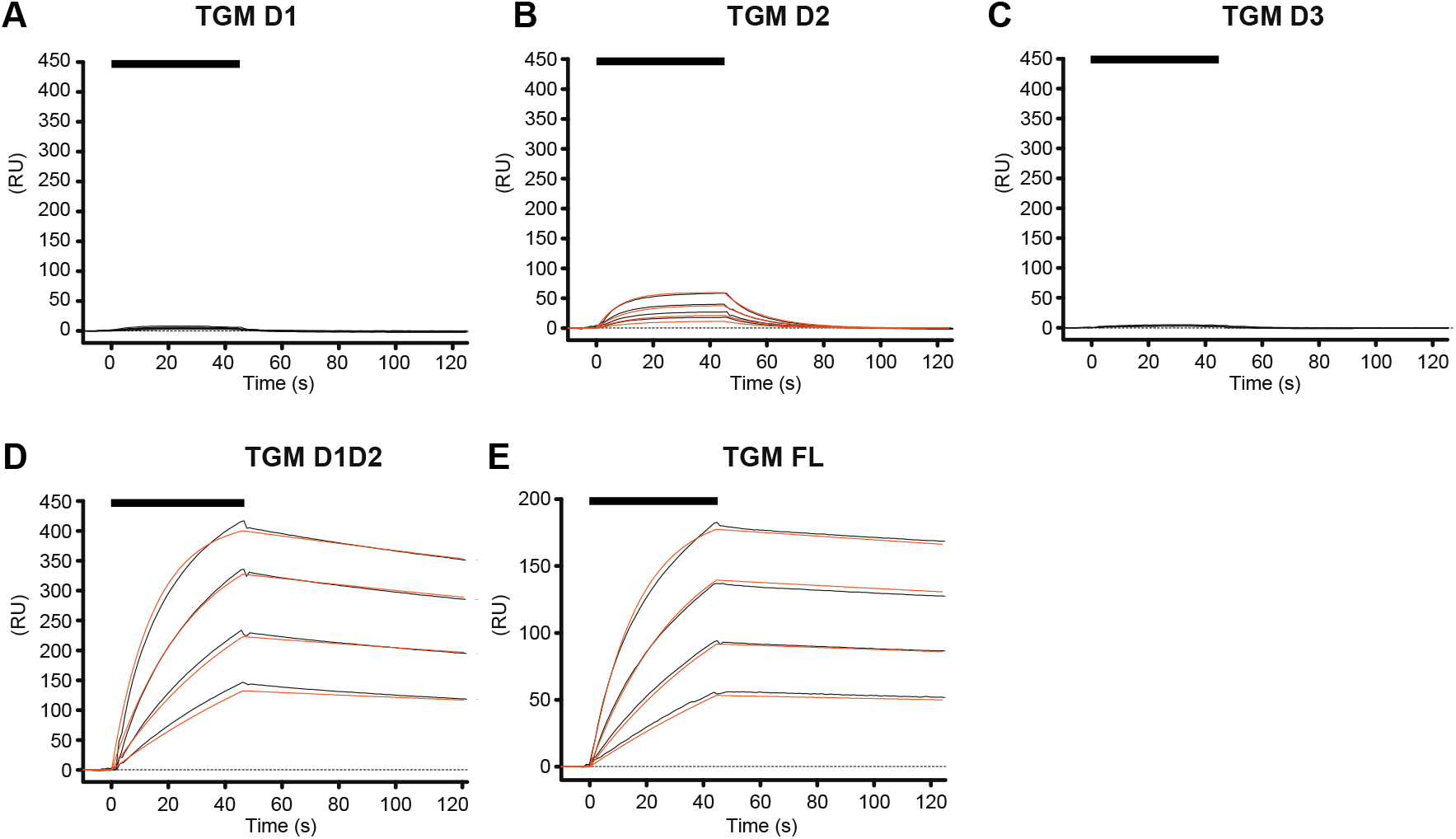
Binding of TβRI by TGM-D1, TGM-D2, TGM-D3, TGM-D1D2, and TGM-FL as assessed by SPR. A-E. SPR sensorgrams obtained upon injection of TGM-D1 (A), TGM-D2 (B), TGM-D3 (C), TGM-D1D2 (D), or TGM-FL (E) over immobilized TβRI. Sensorgrams, obtained upon injection of a two-fold dilution series of each TGM construct (TGM-D1 0.031 – 1.0 μM, TGM-D2 0.125 – 1.0 μM, TGM-D3 0.125 – 1.0 μM, TGM-D1D2 0.125 – 1.0 μM, and TGM-FL 0.125 – 1.0 μM) are shown in black, with the fitted curves in orange (data for TGM-D1 and TGM-D3 were not fit due to weak signal). Black bars shown in the upper left specify the injection period.

### T/JRII utilizes a similar set of residues to bind TGM-D3 and TGF-β

To determine if TβRII might bind to TGM-D3 with some of the same residues that it uses to bind TGF-β3, ITC competition binding experiments were performed. The TGF-β3/TGM-D3 competition experiments with TβRII were performed using an engineered TGF-β monomer, known as mmTGF-β2-7M2R, rather than TGF-β3. mmTGF-β2-7M2R has an intact finger region and binds TβRII with the same affinity as TGF-β1 and TGF-β3^39^, but unlike TGF-β1 or TGF-β3, is highly soluble at neutral pH. The ITC competition measurements were performed by titrating mmTGF-β2-7M2R into the sample cell loaded with TβRII in either the absence or presence of increasing concentrations of TGM-D3 (Fig. 6A-C). The addition of TGM-D3 both increased the extent of curvature in the binding isotherms and reduced the overall enthalpy, consistent with the behavior expected for competitive binding. To quantify this, the integrated heat from the three experiments, together with fitted K_D_ and enthalpy for the TGM-D3:TβRII interaction, were globally fit to a simple competitive binding model to derive the binding constant for the high affinity mmTGF-β2-7M2R:TβRII interaction in the absence of competitor (Fig. 6D-F, Table 3). The K_D_ for the high affinity mmTGF-β2-7M2R:TβRII interaction was found to be 35.20 nM (17.16, 64.42 – 68.3% CI), in accord with previous SPR measurements for the TβRII:TGF-β interaction with immobilized TGF-β1 or TGF-β3^39^. The clear evidence of competition demonstrated in this experiment shows that TβRII uses some or all of the same residues to bind TGF-β and TGM-D3.

**Figure 6.**
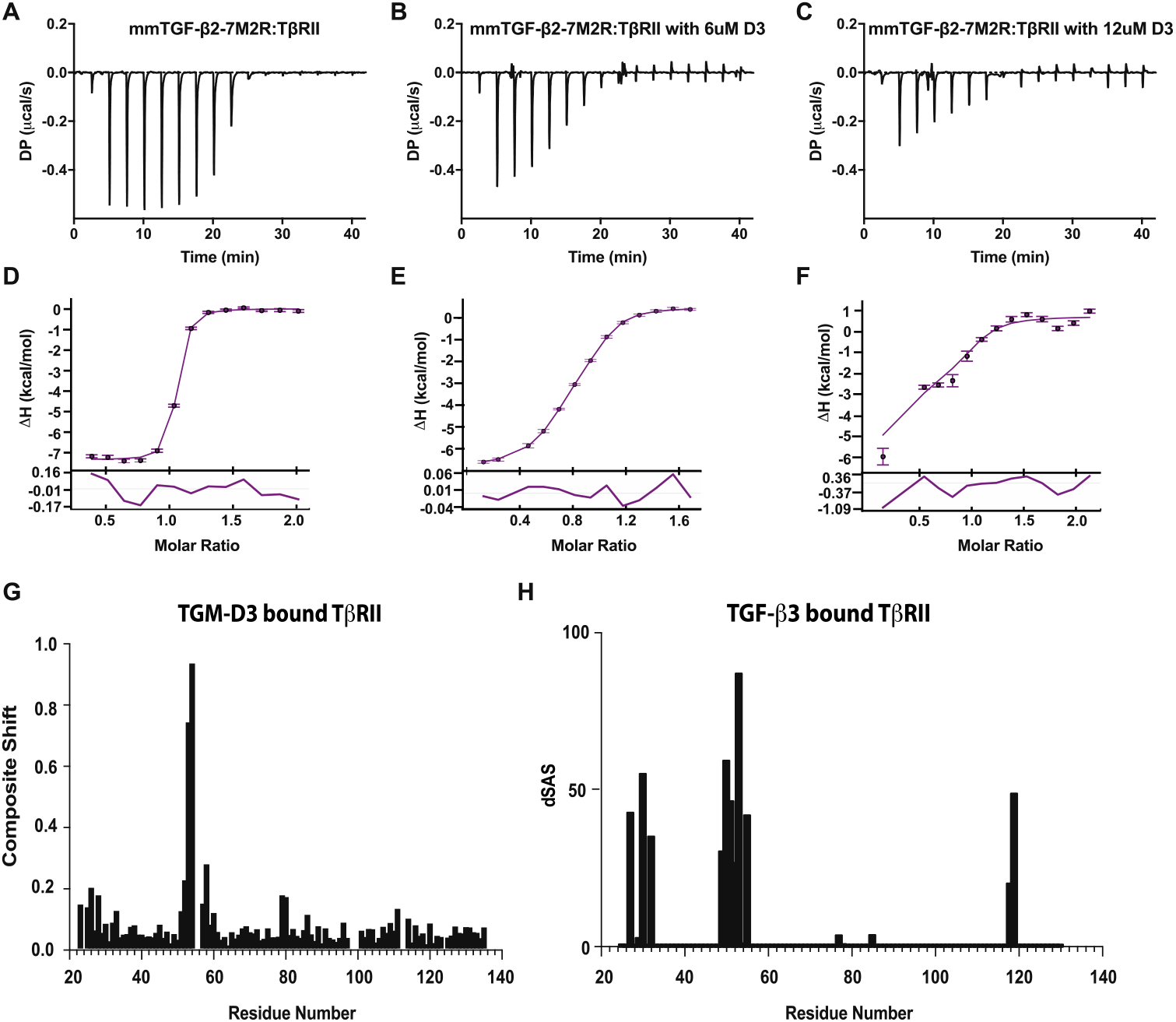
TGM-D3 and mmTGF-β2 7M2R competitive binding to TβRII. A-C. ITC thermograms for injection of 150 μM mmTGF-β2 7M2R into 15μM TβRII in the sample cell with 0 μM (A), 6.0 μM (B), or 12.0 μM (C) TGM-D3. All experiments were performed at 35 °C with a sample cell and syringe buffer consisting of 25 mM sodium phosphate, 50 mM NaCl, pH 6.0. D-F. Global fit of the integrated heats with residuals below global fit (0 mM TGM-D3 (D), 6 μM TGM-D3 (E), and 12 μM TGM-D3 (F)) from the experiments shown in panels A-C to a competition binding model in which both TGM-D3 and mmTGF-β2 7M2R form 1:1 complexes with TβRII. G-H. Composite shifts of TβRII upon binding to TGM-D3 and the change in solvent accessible surface areas of TβRII alone relative to that bound to TGF-β3 G. Composite shift perturbations determined through normalization of chemical shift perturbations for each type of nucleus analyzed between bound and unbound TβRII and plotted for each individual residue. H. Difference in solvent accessible surface (Å^2^) area plotted for individual residues of TβRII between free and TGF-β3 bound form using PDB 1KTZ.

**Table 3.** ITC-based TβRII competition binding.

|  |  |
| --- | --- |
| Cell | TβRII |
| Syringe | mmTGF-β7M-2P |
| Competitor<sup>1</sup> | 0, 6.0, or 12.0 μM TGM-D3 |
| Cell concentration (μM) | 15 |
| Syringe concentration (μM) | 150 |
| Temperature (°C) | 35 |
| K<sub>D</sub><sup>2</sup> (nM) | 35.20 (17.16, 64.42) |
| ΔH<sup>2</sup> (kcal mol<sup>-1</sup>) | -7.36 (-7.70, -7.03) |
| ΔG<sup>3</sup> (kcal mol<sup>-1</sup>) | -10.51 |
| -TΔS<sup>3</sup>(kcal mol<sup>-1</sup>) | -3.15 |

<sup>1</sup>Competitor was added to the sample cell
<sup>2</sup>K<sub>D</sub> and ΔH correspond to the parameters, derived from the global fit, for TβRII:mmTGF-β binding in the absence of competitor, uncertainty determined by 68.3% confidence interval
<sup>3</sup>ΔG and -TΔS correspond to those for TβRII:mmTGF-β binding in the absence of competitor calculated from ΔG = ΔH- TΔS and globally fitted values for K<sub>D</sub> and ΔH

To investigate this further, the backbone of ^15^N, ^13^C TβRII was fully assigned as bound to unlabeled TGM-D3. The assigned chemical shifts for the bound form were then compared to those previously reported for the unbound form^40^ (Fig. S9). The largest shifts, as assessed from a composite of the backbone N, Ca, CO, and sidechain Cβ chemical shift perturbations (CSPs), fell within a narrow region from residue 52-54 (Fig. 6G). This pattern of CSPs was compared to the change in solvent accessible surface area (SAS) of TβRII upon binding TGF-β. The regions of TβRII most hidden by solvent upon binding TGF-β fell within a similar area from residue 50-56 (Fig. 6H). This commonality between the SAS of TβRII bound by TGF-β and the CSPs of TβRII bound by TGM-D3 confirms that TβRII uses the same primary motif, an edge β-strand that binds deeply in the cleft between the fingers 1-2 and 3-4^40,41^, to bind both ligands. TGM-D3 leads to only minor shift perturbations outside of the edge β-strand noted above, whereas TGF-β3 decreases solvent accessibility in regions that flank this strand, suggesting that TGF-β3 has more extensive contacts with TβRII than TGM-D3, an observation consistent with TβRII’s considerably higher affinity for TGF-β1/-β3 compared to TGM-D3 (K_D_s ca. 30-50 nM and 500-1000 nM, respectively).

### T/JRI utilizes a similar set of residues to bind TGM-D2 and TGF-β:TβRII

To determine if TβRI might bind to TGM-D1D2 with some of the same residues that it uses to bind the TGF-β3/TβRII complex, ITC competition binding experiments were performed by titrating TGM-D1D2 into the sample cell loaded with TβRI in either the absence or presence of saturating concentrations of TGF-β3/TβRII. The control titration of TGF-β:TβRII into TβRI alone demonstrated a fitted K_D_ of 61.7nM (35.7 – 97.2nM, 68.3% CI) (Fig. 7A-B), which is similar to that for TGM-D1D2 binding TβRI alone (K_D_: 25.3nM (10.7 – 48.3, 68.3% CI)).

**Figure 7.**
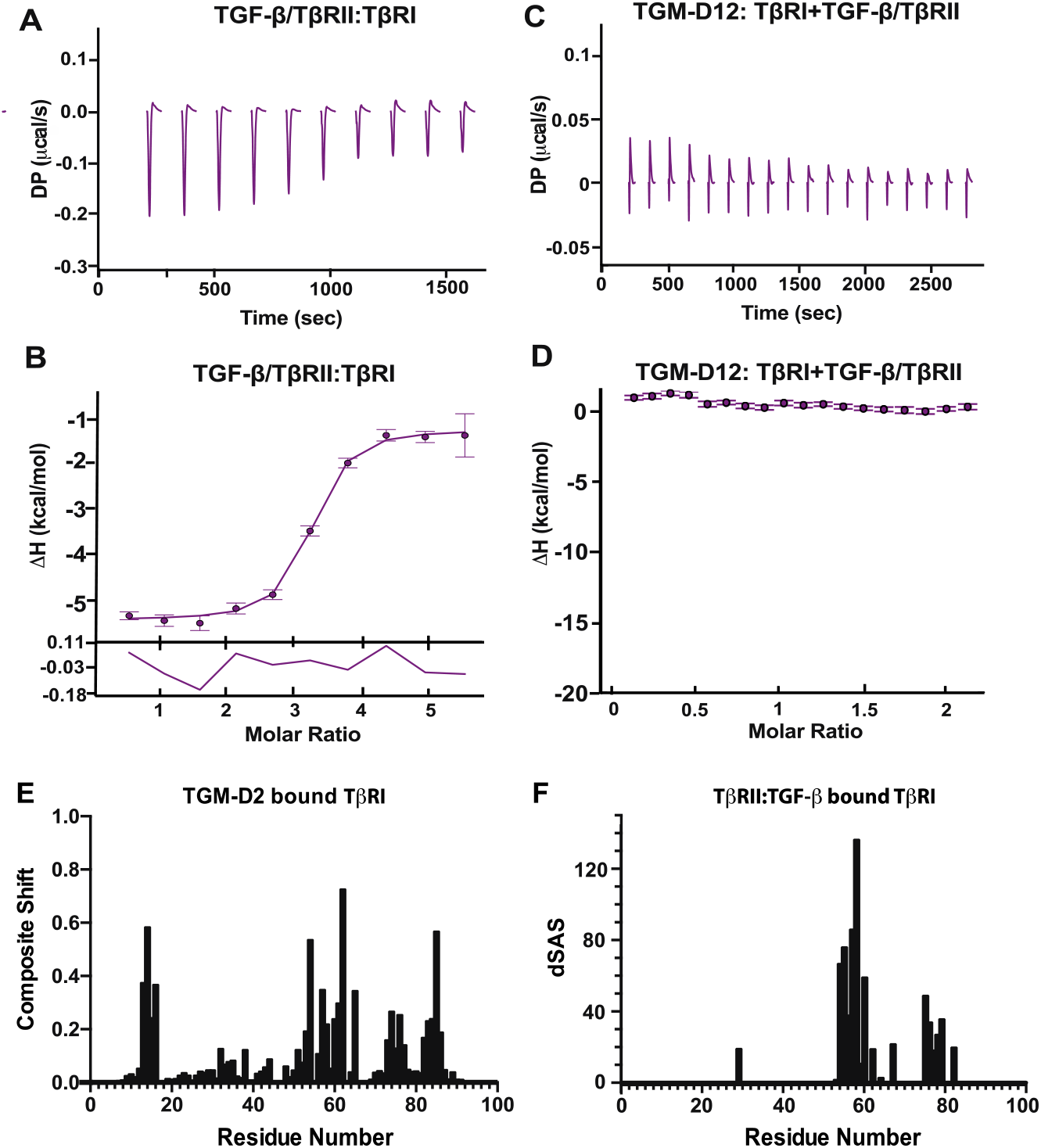
TGM-D1D2 and TGF-β3/TβRII competitive binding to TβRI. A, C. Thermograms obtained upon injection of 1:2 TGF-β/TβRII (A) or TGM-D12 into TβRI with a saturating concentration of TGF-β/TβRII binary complex (E). B, D. Integrated heats for the titrations shown in panels A and C with residuals. The data points correspond to the integrated heats and the purple lines correspond to a fit to the model for 1:1 binding (B,D). All experiments were performed at 25 °C with a sample cell and syringe buffer consisting of 25 mM HEPES, 50 mM NaCl, 0.05% NaN_3_ pH 7.5. E-F. Composite shifts of TβRI binding to TGM-D2 and the change in the solvent accessible surface area (SAS) of TβRI upon binding TGF-β/TβRII (E). Composite shift perturbations for each type of nucleus analyzed between bound and unbound TβRI and plotted for each individual residue. Composite shifts were determined through normalization of chemical shift perturbations between bound and unbound TβRI and plotted for each individual residue. (F). Difference in SAS plotted for individual residues of TβRI between free and bound TGF-β3:TβRII bound, using PDB 2PJY.

However, unlike the TGM-D1D2:TβRI interaction which had a large enthalpy, −18.70 kcal mol^-1^, the TGF-β:TβRII:TβRI interaction had a much smaller enthalpy −4.21 (−4.49 – −3.95) kcal mol^-1^, even at increased temperature (Table 4), indicating that the binding reaction is more entropically driven. Owing to similar K_D_s but a significantly greater enthalpy for the TGM-D1D2:TβRI interaction, the competition experiment was performed by saturating TβRI in the cell with TGF-β:TβRII binary complex and then titrating TGM-D1D2 using the syringe (Fig. 7C-D). The resultant titration yielded minimal heat relative to the TGM-D1D2:TβRI interaction (Fig. 4D, G) and could not be quantitively fit, indicating that the TGF-β:TβRII binds to some or all of the same residues of TβRI at TGM-D1D2.

**Table 4.** ITC-based TβRI competition binding.

|  |  |  |
| --- | --- | --- |
| Cell | TβRI | TβRI + TGF-β/TβRII |
| Syringe | TGF-β/TβRII | TGM-D1D2 |
| Competitor | 0 | 6<sup>2</sup> |
| Cell concentration (μM) | 5<sup>1</sup> | 10 |
| Syringe concentration (μM) | 100 | 110 |
| Temperature (°C) | 30 | 25 |
| K<sub>D</sub> (nM) | 61.4 (35.7-97.2) | NA<sup>3</sup> |
| ΔH (kcal mol<sup>-1</sup>) | -4.21 (-4.49 – -3.95) | NA<sup>3</sup> |
| ΔG (kcal mol<sup>-1</sup>) | -10.00 | NA<sup>3</sup> |
| -TΔS (kcal mol<sup>-1</sup>) | -5.79 | NA<sup>3</sup> |

<sup>1</sup>Competitor was added to the syringe in a concentration of 5μM dimer with n=2
<sup>2</sup>Competitor was added to the sample cell in a concentration of 6μM dimer with n=2
<sup>3</sup>K<sub>D</sub>, ΔH, ΔG and -TΔS were unable to be fitted

To investigate this further, the backbone of ^15^N, ^13^C TβRI was fully assigned as bound to unlabeled TGM-D2 (Fig. S10). TGM-D1D2 was not used due to its higher molecular weight and reduced solubility in TβRI-compatible NMR buffers. The assigned chemical shifts for the bound form were then compared to those previously reported for the unbound form^17^. The largest shifts, as assessed from a composite of the backbone N, Ca, CO, and sidechain Cβ CSPs, fell within four distinct regions: 1) residues 13-16, 2) residues 53-65, 3) residues 73-77, and 4) residues 82-86 (Fig. 7E). This pattern of shifts was compared to the change in SAS of TβRI upon binding the TGF-β/TβRII complex. The regions of TβRI protected from solvent upon binding TGF-β and TβRII fell within similar areas from residue 54-60 and 75-78 (Fig. 7F). The commonality between the change in SAS of TβRI upon binding TGF-β:TβRII and the CSPs of TβRI upon binding TGM-D2 confirms that TβRI uses the same primary motifs, the pre-helix extension (residues 53-65) that bridges β-strands 4 and 5 and the C-terminal end of β-strand 5 (residues 73-77), to bind both ligands. TβRI has two regions of TGM-D3-inncuded CSPs outside these regions, residues 13-16 and 82-85; these residues are positioned adjacent to the C-terminal end of β-strand 5, suggesting that TGM-D2 has a larger, more distributed contact surface with TβRI as compared to TGF-β:TβRII.

### TGM-D3 structure and dynamics

The structure of TGM-D3 was determined based on ^1^H-^1^H NOE distance restraints, ^1^H-^15^N, ^13^C^a^-^1^H^a^, and ^13^C^O^-^15^N RDCs, and ^3^J^HN-Ha^, 3J^Ha-Hβ^, 3J^HN-Hβ^ J-couplings and essentially complete chemical shift assignments for both the backbone and sidechains (Table 5). The backbone root-mean-square deviation (RMSD) for the ten lowest-energy structures relative to the lowest energy structure was 0.41 Å when aligned according to the regions of regular secondary structure, or 2.31 Å when aligned over the length of the ordered core (Fig. 8A, Table 5). TGM-D3 is comprised of four beta strands (Val^15^-Gly^21^, Thr^45^-Cys^51^, Glu^62^-Lys^69^, Ser^76^-Tyr^80^) arranged into a highly twisted antiparallel β-sheet with a β1:β2:β3:β4 topology (Fig. 8A). There is also a 3_10_helix (Gln^56^-Ala^58^) connecting β2 and β3 in some, but not all of the lowest-energy structures (Fig. 8A). The structures are consistent with that derived from an analysis of secondary shifts^42^, with four high probability extended regions predicted between residues 12-19, 44-50, 62-69, and 76-80, and a low probability helical region from residues 54-56 (Fig. S11C). The secondary shifts also predict, with lower probability, extended regions between residues 5-7 and 29-34. The former corresponds to the N-terminal region, while the latter corresponds to the middle section of the 23-residue hypervariable (HVL) loop that connects β1 and β2 (Fig. 8A). This section of the HVL from extends perpendicularly across the C-terminal end of β1 and is mostly converged among the ten lowest energy structures, with an average pairwise RMSD of 0.85 Å. The segments from residues 5-7 and 29-34, although highly extended, do not form hydrogen bonds that define a β-strand and thus are not classified as such in the calculated structures.

**Table 5.** TGM-D3 Structural Statistics.

|  |  |
| --- | --- |
| **NOE** |  |
| Intramolecular NOE: $i-j = 0$ | 465 |
| Sequential NOE: $i-j = 1$ | 323 |
| Short-Range NOE: $1 < i-j < 5$ | 104 |
| Long-Range: $i-j \geq 5$ | 247 |
| **Angle** |  |
| TALOS ($\phi, \psi$) dihedral constraints | 136 |
| $\chi_1$ Angle | 12 |
| $^3J_{\text{HNHA}\alpha}$ | 44 |
| **RDC** |  |
| RDC: N-H | 69 |
| RDC: $\text{H}\alpha\text{-C}\alpha$ | 74 |
| RDC: $\text{C}\alpha\text{-CO}$ | 66 |
| **RMSD (Deviations)** |  |
| Bonds (Å) | $0.008 \pm 0.000$ |
| Impropers (°) | $1.195 \pm 0.135$ |
| Angles (°) | $1.160 \pm 0.085$ |
| Dihedral (°) | $5.041 \pm 0.920$ |
| HBDA (Å) | $0.021 \pm 0.015$ |
| $^3J_{\text{HNHA}\alpha}$ (Hz) | $1.349 \pm 0.0193$ |
| **Ramachandran<sup>a</sup>** |  |
| Most Favored | 81.2% |
| Additionally Allowed | 14.5% |
| Generously Allowed | 2.9% |
| Disallowed | 1.4% |
| **RMSD<sup>b</sup>** |  |
| <u>Secondary Structure<sup>c</sup></u> |  |
| Backbone | 0.41 Å |
| Heavy | 1.87 Å |
| <u>Core<sup>d</sup></u> |  |
| Backbone | 2.31 Å |
| Heavy | 3.76 Å |

<sup>a</sup>Ramachandran plot for the lowest-energy structure<sup>b</sup>RMSD values are the average values of the pairwise RMSD comparisons of the top 10 structures<sup>c</sup>Residues 15-21, 45-51, 56-58, 62-69, 76-80<sup>d</sup>Residues 6-81

**Figure 8.**
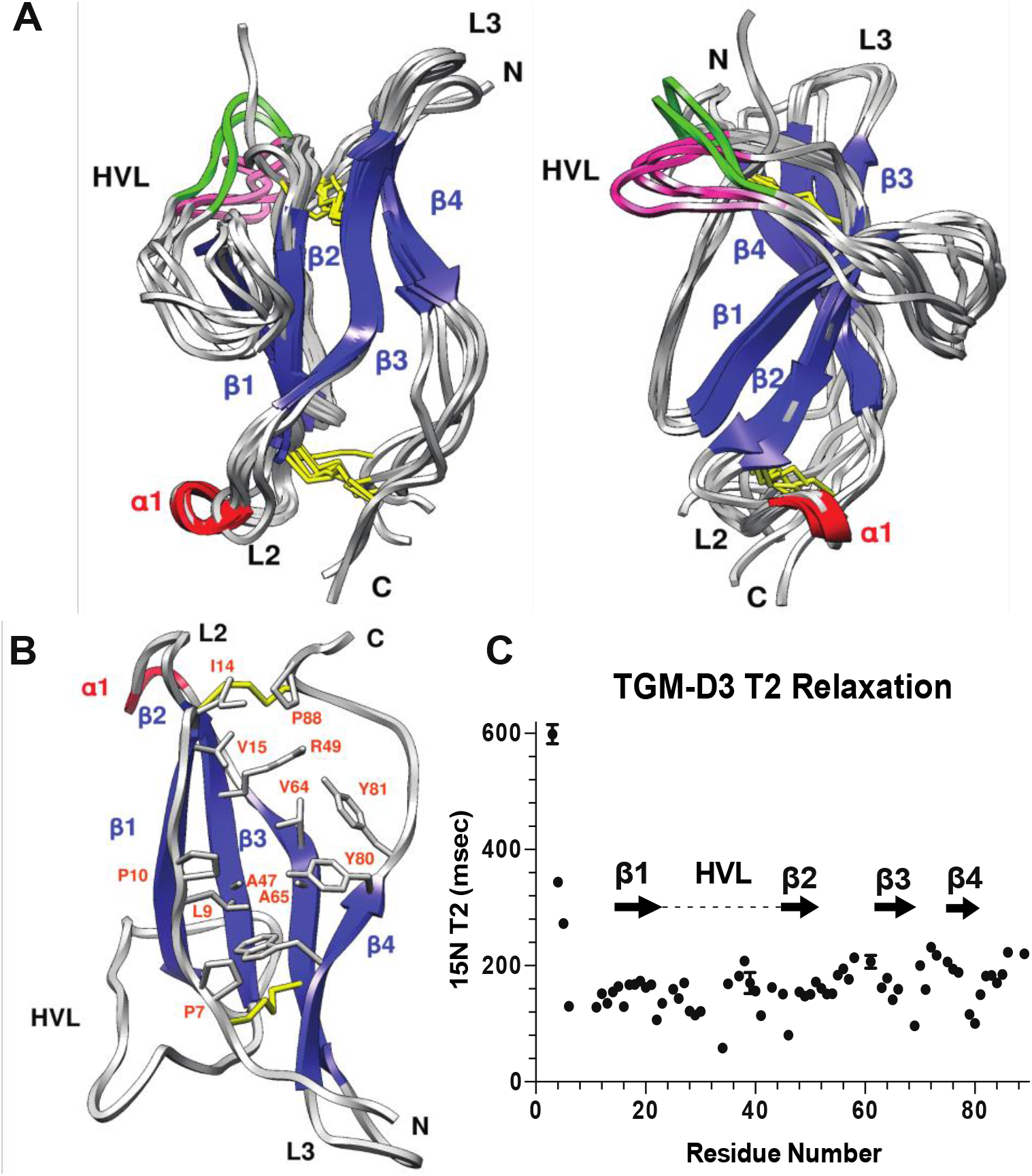
NMR structure of TGM-D3: Dynamics and Structural Properties. A. Ensemble of the five-lowest energy NMR structures of the unbound form of TGM-D3 as determined by Xplor-NIH after refinement: β-strands, dark blue; loops, gray; 3_10_ helix, red; disulfide bonds, yellow, multiple conformations of HVL highlighted in magenta and green. Key structural features are indicated. B. Representative structure of TGM-D3; rotated 180° around the x-axis in comparison to A. Side-chains of hydrophobic core residues presented and labeled in orange. β-strands, dark blue; loops, gray; 3_10_ helix, red; disulfide bonds, yellow Key structural features are indicated. C. Backbone ^15^N T_2_ Relaxation time for TGM-D3 plotted per individual residue with structural features mapped (β-strands, arrows; HVL, dashed line).

The Cys^6^-Cys^67^ disulfide pins the N-terminus to one end of the concave surface of the β-sheet, while the C-terminus is pinned to the other end of the sheet by the Cys^51^-Cys^87^ disulfide (Fig. 8B). This creates a large broad cavity that is bordered on one edge by the extended N-terminal segment from Cys^6^-Gly^13^ and on the other by β4 and the extended segment that follows, which spans from Ser^76^-Cys^87^ (Fig. 8B). The core of the protein is formed by hydrophobic residues in the cavity, and includes Leu^9^ and Ile^14^ from the extended N-terminal segment, Val^15^ and Tyr^17^ from β1, Ala^47^ and the hydrophobic portion of the sidechain of Arg^49^ from β2, Val^64^ and Ala^65^ from β3, and Trp^78^, Tyr^80^, and Tyr^81^ from β4 (Fig. 8B). Though the majority of these residues are completely buried, including Leu^9^, Ala^47^, Val^64^, Ala^65^, and Trp^78^, there are a few that are partlly exposed, particularly Ile^14^ and Val^15^, both of which are found near the end of the cavity closest to the Cys51-Cys87 disulfide bond where it is at its widest.

The backbone ^15^N T_2_ relaxation times, which are sensitive to both fast (ns-ps) timescale motions that result from low amplitude fluctuations of the backbone, but also to larger amplitude rearrangements that occur on slower (μs – ms) timescales, are significantly increased in the N-terminal tail and modestly increased near the C-terminal end of the HVL and in the shorter loops connecting β2-β3 and β3-β4 (Fig. 8C). The increases in ^15^N T_2_ indicate increased flexibility in these regions, especially the N-terminal tail which does not converge in the final ensemble of structures. The other loop regions converge reasonably well, consistent with their more modest increases in ^15^N T_2_ (Fig. 8A), although one exception is the HVL, which adopts two conformations, in which the C-terminal portion of the HVL either ascends or descends as it contacts the extended N-terminus (Fig. 8A, green and pink respectively).

### TGM-D3 compared to other CCP domains

CCP domain structures with the closest fold to TGM-D3, as identified by a DALI^43,44^ search of the protein data bank, have close correspondence in the four β-strands that form the core of the CCP fold, but have two additional β-strands, one in the loop connecting β2 and β3, designated β’, and another at the C-terminus, designated β’’ (Fig. 9A). The β’ and β’’ strands are present in all of the top-scoring CCP domains and pair with one another (Fig. 9B). This serves to draw the C-terminal segment toward the loop connecting β2-β3 and essentially eliminates the large broad cavity that is formed between the extended N-terminus and β4 (Fig. 9C). Thus, unlike TGM-D3, the hydrophobic core of other CCP domains, which also includes a conserved tryptophan, is entirely buried and there are no partly exposed hydrophobic residues Additionally, none of these CCP domains, nor any other known CCP domain, contains the long HVL extension similar to the one in TGM-D3, which as noted is largely ordered and wraps laterally around the CCP domain on the convex surface of the sheet. Instead, the HVL in other CCP domains simply extends into solvent to connect β1 and β2 and has no contact with the convex surface of the sheet.

**Figure 9.**
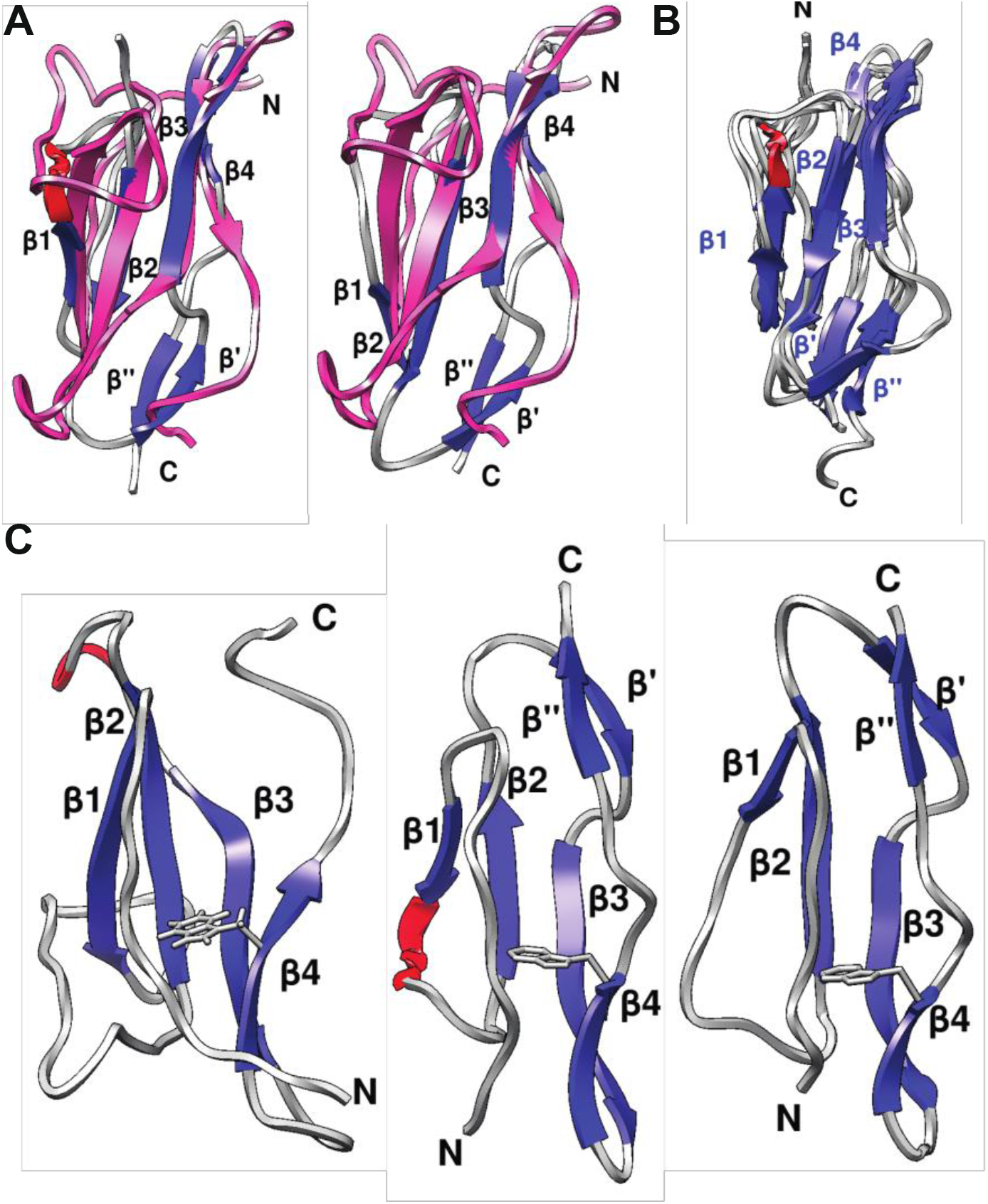
TGM-D3 comparison to CCP domains. A. Alignment of TGM-D3 (pink) to representative CCP domain (1ckl left, 1h2p right: β-strands, dark blue; loops, gray; 3_10_ helix, red. Key structural features are indicated. B. Ensemble of CCP domains (PDB: 2psm, 1ckl, 1h2p, 5fo9, 5foa) aligned using UCSF Chimera: β-strands, dark blue; loops, gray; 3_10_ helix, red. C. Representative structure of hydrophobic core of TGM-D3 (left), CD46 (center, PDB: 1ckl), CD55 (right, PDB: 1h2p). β-strands, dark blue; loops, gray; 3_10_ helix, red. Key structural features are indicated. Conserved tryptophan residue shown in all three structures.

### TGM-D3 engages its binding partner T/JRII in a manner distinct to that of the canonical CCP domain

To determine the TGM-D3:TβRII binding interface, the backbone of ^15^N, ^13^C TGM-D3 was fully assigned as bound to unlabeled TβRII (Fig. S11A-B). The two regions that were most strongly perturbed include residues 62-71 and 77-85, which correspond to most of β3 and β4, as well as a few residues that extend beyond the end of β4 (Fig. 10A). The regions that are perturbed to a lesser extent include residues 42-47 and 21-28, which correspond to the N-terminal end of β2 and the N-terminal end of the HVL as it emerges out of β1 and makes a sharp turn before extending laterally across the C-terminal end of β1. The residues that are most strongly perturbed, including Phe^63^, Ile^66^, Tyr^80^, Tyr^81^, and Ile^84^, lie within the large cavity on the concave face of TGM-D3 (Fig. 8B, 10A). The largest shift perturbations in TβRII, as previously noted, are from residue 52-54 (Fig. 6G), which corresponds to an edge β-strand. Through previous crystal structures of TβRII bound to TGF-β^17,45,46^, TβRII has been shown to insert this edge β-strand into the hydrophobic pocket formed between the fingers 1-2 and 3-4 of TGF-β. The region of TGM-D3 that is perturbed upon binding TβRII is similarly hydrophobic (Fig. 8B, 10A), and thus could conceivably accommodate the edge β-strand of TβRII.

**Figure 10.**
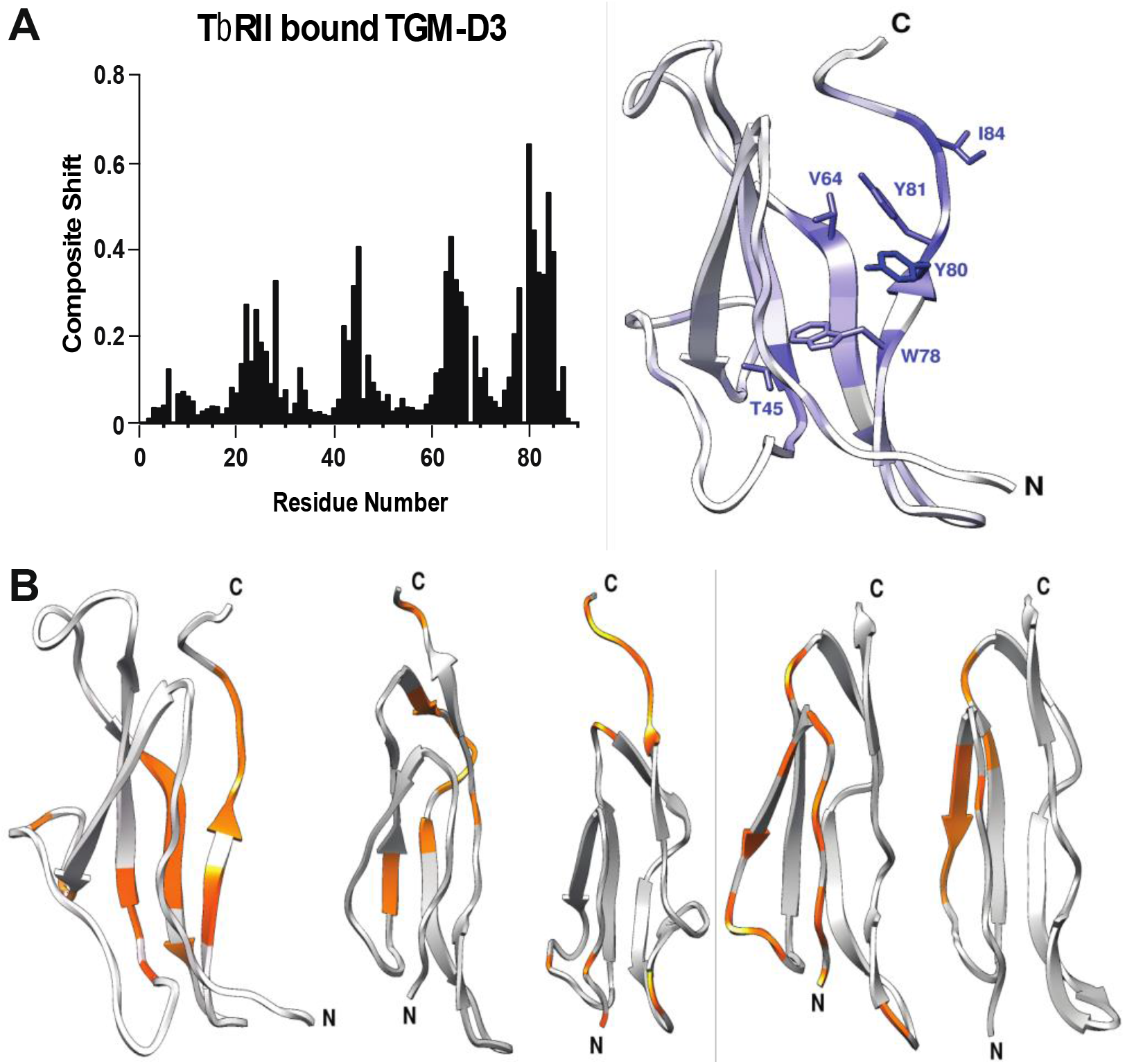
TGM-D3 binding interface comparison to CCP domain binding interface. A. Composite shift perturbations determined through normalization of chemical shift perturbations for each type of nucleus analyzed between bound and unbound TGM-D3 and plotted for each individual residue (left). TGM-D3 colored using a scale where white indicates minimal shift perturbation and blue indicates maximal perturbation (right). B. Structures of TGM-D3 (left) and CCP domains (IL-15, PDB: 2psm. MASP, PDB: 4fxg. CR1, PDB: 5fo9. CD55, PDB: 6la5 – in order from left to right) with amino acids involved in binding surfaces colored by threshold for composite shift > 0.2, (TGM-D3) or by change in solvent accessible surface area > 20Å^2^ (CCP domains).

The binding surface that TGM-D3 uses to bind TβRII is distinct relative to that used by other CCP domains to bind their partners. Though there is no single interface that defines how CCP domains bind their partners, due to both their ubiquity and the range of partners they bind, an analysis of those CCP domains most closely related to TGM nonetheless shows that they generally use their convex surface, specifically the extended N-terminal segment, β1, β2, and the β2-β3 loop to bind their ligands (Fig 10β). Thus, TGM-D3 has evidently acquired two large structural modifications that engender it with an alternate manner of binding its partner TβRII: (1) the long, laterally protruding, and structurally-ordered HVL of TGM-D3 likely acts to block binding to the convex face of TGM-D3, thus preventing interactions with other partners through the interface that is commonly employed, and (2) the large broad cavity between the extended N-terminus and β4, which is formed due to the elimination of the β’ and β’’ strands, and replacement of the former with a 3_10_ helix, accommodates the edge b-strand of TβRII in a manner that mimics binding between the fingers of the TGF-β in the TGF-β:TβRII complex.

## Discussion

The genome of the mouse helminth *H. polygyrus* encodes a highly expanded family of CCP-containing proteins, several of which have been identified in its secretome to regulate host immune response. This distinguishes *H. polygyrus* uniquely from closely related helminths, such as the sheep hookworm *Teladorsagia circumcincta* and the human hookworm *Necator americanus* which do not have an expanded CCP family and attenuate host immune response through other mechaniisms^47-50^. TGM and its five adult (TGM-2 through TGM-6) and four larval (TGM-7 through TGM-10) homologues are amongst the proteins in this expanded family, and as noted, at least four of these, TGM, TGM2, 3, and 4, regulate immunosuppressive signaling through the T_reg_ pathway. Though potency of signaling through TGM is similar to that of TGF-β^12^, protein-protein binding kinetics and amplitude of signaling in reporter cell lines is distinct, as is the overall gene expression profile, with increased T_reg_ potency, and decreased fibrotic gene response^12^. HpARI and HpβARI, which impact innate immune responses, are also among the CCP-containing proteins encoded by *H. polygyrus*, although the sequences of the HpARI and HpβARI CCPs are highly divergent from those of TGM and its homologs.

The results presented demonstrate that TGM binds the TGF-β receptors, TβRI and TβRII, in a modular manner, with TGM-D2 and TGM-D3 the main partners, respectively. The binding of TβRI, is however significantly potentiated by TGM-D1. The underlying basis for this potentiation is likely binding of TβRI through a composite TGM-D1:TGM-D2 interface as the NMR titration data presented in Fig. S8B shows that TGM-D1 directly, albeit weakly, binds TβRI. This manner of binding, however, is unsurprising given that it is common for CCP-containing proteins to bind partners through arrays of CCPs, with avidity playing an important role, and with the CCP domains generally connected by short linkers, as is the case for TGM domains 1 and 2.

The overall manner by which TGM binds TβRI and TβRII and assembles them into a signaling heterodimer is distinct in comparison to the TGF-β homodimers, which assemble a (TβRI:TβRII)_2_ heterotetramer, first by binding TβRII with moderate to high affinity (K_D_ ca. 50 nM), and in turn recruiting and binding TβRI through a composite TGF-β:TβRII interface with high affinity (K_D_ ca. 30 nM). Though further studies are required, the differences in stoichiometry of the TGM vs. TGF-β signaling complex, the differences in the manner of assembly, as well as potential differences in the overall architecture of the signaling complex, specifically how the type I and type II kinases are arrayed relative to one another in the cytoplasm, likely explains the differences in the amplitude and kinetics of signaling and overall shift of the gene expression profile away from extracellular matrix accumulation and towards immunosuppression.

The ITC competition binding and NMR assignments of the free and bound forms of TβRI and TβRII clearly demonstrate that TGM-D1D2 and TGM-D3 truly mimic the mammalian cytokine by engaging the same primary motifs of the receptors, the -PRDRP-pre-helix extension in TβRI and the β4 edge β-strand in TβRII. The structure of TGM-D3, together with identification of the binding site for TβRII based on NMR assignments of the free and TβRII-bound forms, provides a remarkable demonstration of how TGM-D3 has adapted, relative to other CCP domains, to uniquely and specifically bind TβRII through the cavity formed on its concave surface.

Though all of the domains of TGM are predicted to have the overall fold of CCP domain, only TGM-D3 binds to TβRII. Sequence comparisons of TGM-D3 with the other domains of TGM (Fig. S12A), demonstrates that all of the domains contain the HVL insertion and two disulfide bonds. TGM-D3 is however unique in that the β3-β4 loop is 5-6 residues shorter in the other domains compared to domain 3, thus this loop is likely a tight β-turn rather than a more extended turn as in domain 3. This may potentially alter the overall shape and dimensions of the cavity on the concave surface to accommodate other receptors of the TGF-β family, all of which, including TβRI, have been shown to engage their cognate ligands through structural motifs distinct from the β4 edge strand of TβRII.

Of the TGM family members in the adult parasitic secretome, TGM, TGM-2,-3,-4, and −6 have each been shown to bind TβRII^29^ (TGM-4 and TGM-6 data unpublished), consistent with sequence alignments which show high conservation amongst the our β-strands, the loop connecting β2 and β3, and the extended HVL, with minor amino acid differences that likely do not correlate to larger structural differences (Fig. S12B). Thus, it is likely that domain 3 of each of these homologs have a large cavity bordered by the extended N-terminal segment and β4 that binds TβRII, while binding of other potential partners through the convex face is prevented by the extended HVL.

This specific CCP domain modification likely does not extend to other CCP proteins in the *H. polygyrus* excretory-secretory products (HES). HpARI, and HpβARI, which are involved in binding Il-33 receptor ST-2, are predicted to be somewhat structurally dissimilar to TGM-D3. HpARI has three CCP domains, and HpβARI has two CCP^31-33^, and analysis of the secondary structure of these using the PHYRE2 server^51^, reveals significant differences: HpARI-CCP1 is predicted to have 5 β-strands, CCP2 is predicted to have 8 β-strands, together with two predicted helices, and CCP3 is predicted to have 4 β-strand regions and a large helical region (Fig. S13A). HpβARI-CCP1 and CCP2 are both predicted to have six β-strands, which likely means that these two proteins have the β’ and β’’-strands that TGM-D3 lacks, and likely lack a cavity on the concave surface (Fig. S13B). Thus, the unique modifications of TGM-D3, and likely other TGM domains as well, are not likely shared by other by other CCP-containing in the *H. polygyrus* secretome, indicating that these modifications have arisen to inure these domains with the remarkable ability to specifically bind the type I and type II receptors of the TGF-β family.

These findings highlight the unique nature of *H. polygyrus*-mediated immunomodulation. Unlike TGF-β, which binds TβRII and then recruits TβRI to the TGF-β:TβRII binding interface, TGM binds TβRI and TβRII in a modular manner, with TGM-D3 alone binding TβRII and TGM-D1D2 binding TβRI, with higher affinity binding to TβRI than to TβRII. Though the parasite product responsible is structurally dissimilar to TGF-β, TGM has convergently evolved the CCP domain protein family to take advantage of the same type I and type II receptor binding sites as human TGF-β. This structural knowledge can be used both to develop TGM and individual domains of TGM as therapeutic TGF-β mimics and to further explore the role of CCP domain-heavy proteins in parasitic host immunomodulation.

## Materials and Methods

### Expression and purification of TGM domains

DNA fragments corresponding to individual domains of *H. polygyrus* TGM, TGM-D1, TGM-D2, TGM-D3, and TGM-D1D2, were inserted between a KpnI and HindIII sites in modified form of pET32a (EMD-Millipore, Danvers, MA) to include a KpnI site immediately following the coding sequence for the thrombin recognition sequence. The resulting constructs, which included a thioredoxin-hexahistidine tag-thrombin cleavage site-TGM domain coding cassette (Table S1), were overexpressed in βL21(DE3) cells (EMD-Millipore, Danvers, MA) cultured at 37°C. Unlabeled samples for binding studies were produced on rich medium (LB), while ^15^N and ^15^N,^13^C samples for NMR studies were produced using minimal medium (M9) containing 0.1% ^15^NH_4_Cl (Cambridge Isotope Laboratories, Tewksbury, MA) or 0.1% ^15^NH_4_Cl and 0.3% U-^13^C-D-glucose (Cambridge Isotope Laboratories, Tewksbury, MA). Carbenicillin was also included in the media at 50 μg mL^-1^ to select for cells bearing the expression plasmid. Protein expression was induced by adding 0.8 mM IPTG when the light scattering at 600 nm reached 0.75.

Cell pellets from 3 L of culture were resuspended in 100 mL lysis buffer (50 mM Na_2_HPO_4_, 100 mM NaCl, 5 mM imidazole, 10 μM leupeptin, 10 μM pepstatin, 1 mM benzamide, pH 8.0) and sonicated. Following centrifugation (20 min, 15000g), the pellet was washed with 50 mL water, resuspended in 50 mM Na_2_HPO_4_, 100 mM NaCl, 5 mM imidazole, 10

μM leupeptin, 10 μM pepstatin, 1 mM benzamide, 8 M urea, pH 8.0, and stirred overnight at 25 °C. The remaining insoluble material was removed by centrifugation and the supernatant was loaded onto a 50 mL metal affinity column (Ni^++^ loaded chelating sepharose, GE Lifesciences, Piscataway, NJ) pre-equilibrated with 125 mL of resuspension buffer. The column was washed with 100 mL resuspension buffer and the bound protein was eluted by applying a linear gradient of resuspension buffer containing 0.5 M imidazole.

Protein from the eluted peak was treated with reduced glutathione (GSH), such that the final concentration of GSH once the protein was diluted into folding buffer was 2 mM. After a 30 min incubation at 25 °C, the protein was slowly diluted into folding buffer (0.1 M Tris, 1 mM EDTA, 0.5 mM oxidized glutathione (GSSG), pH 8.0) to a final concentration of 0.1 mg mL^-1^ and stirred for 12 - 16 h at 4° C. The folding mixture was concentrated and dialyzed into 25 mM Tris, pH 8.7. Solid thrombin was added to a final concentration of 4 U per milligram of TGM domain and incubated overnight at 25 °C. Cleavage was stopped by the addition of 10 μM leupeptin, 10 μM pepstatin, and 100 μM PMSF, and after re-adjusting the pH to 8.7, the cleavage mixture was passed over a Ni^++^ chelating sepharose column equilibrated with water. Column flow-through, and a subsequent water wash, which contained primarily the TGM domain, was collected. For the TGM-D1 and TGM-D1D2 domains, the flow-through was bound to a Source Q column (GE Lifesciences, Piscataway, NJ) equilibrated in 25 mM CHES, pH 9.0 and eluted with a 0-0.5M NaCl gradient. For the TGM-D2 and TGM-D3 domains, the flow-through was adjusted to pH 5.0 by the addition of acetic acid, bound to a Source S column (GE Lifesciences, Piscataway, NJ) equilibrated in 5 mM sodium acetate, 2M Urea, pH 5.0, and eluted with a 0-0.5M NaCl gradient. Masses of the TGM domains were measured by liquid chromatography electrospray ionization time-of-flight mass spectrometry (LC-ESI-TOF-MS, Bruker Micro TOF, Billerica, MA). TGM-FL was expressed in expi293 cells (Promega, USA) and purified by metal affinity chromatography as previously described^12^.

### Expression and purification of TGF-/J receptor and growth factor constructs

The TGF-B2 mini monomer (mm-TGF-B2-7M2R), TβRII ectodomain, and TβRI ectodomain, were expressed in *E. coli* at 37°C in the form of insoluble inclusion bodies, refolded, and purified as previously described^39,52,53^. Details of the constructs used are provided in Table S1. Masses were verified by LC-ESI-TOF-MS.

### NMR data collection and signal assignments

Samples of TGM-D1, -D2, -D3, and corresponding complexes with TBRI and TBRII, for NMR were prepared at a concentration of 0.03 to 0.2 mM in 25 mM Na_2_HPO_4_, 50 mM NaCl, pH 6.0 and transferred to 5 mm susceptibility-matched microtubes (Shigemi, Allison Park, PA) for data collection. NMR data were collected at 30 °C using a Bruker 600, 700, or 800 MHz spectrometer equipped with a 5 mm ^1^H (^13^C,^15^N) z-gradient “TCI” cryogenically cooled probe (Bruker Biospin, Billerica, MA). Routine ^1^H-^15^N HSQC spectra were acquired with a sequence with sensitivity enhancement^54^, water flipback pulses^55^, and WATERGATE water suppression pulses^56^. Backbone resonances were assigned by recording and analyzing ^1^H-^15^N HSQC and HNCACB, CBCA(CO)NH, HNCA, HN(CO)CA, HNCO, and HN(CA)CO triple resonance data sets. Proton and sidechain resonances were assigned by recording and analyzing ^1^H-^13^C CT-HSQC, CC(CO)NH, HBHACONH, HCCH-TOCSY, H(CC, CO)NH, HNHA, and HNHB data sets. NMR data were processed using nmrPipe^57^ and analyzed using a combination of NMRFAM-SPARKY and the Bayesian-based PINE software packages^58,59^. T_2_ Relaxation experiments were performed with ^15^N TGM-D3 using ZZ-exchange experiments^60^.

### NMR structure determination of TGM-D3

The structure of TGM-D3 was initially calculated using the assigned chemical shifts and measured ^1^H-^15^N residual dipolar couplings (RDCs) as input. Residual dipolar couplings for the backbone amides were measured using an IPAP-HSQC sequence^61^ and an oriented sample containing 10 mg mL^-1^ Pf1 phage. Refined structure of TGM-D3 was determined using the program NIH-XPLOR^62^, with manually peak picked 3D ^15^N-edited and ^13^C-edited NOESY data, backbone ^1^H-^15^N ^1^Ha-^13^Ca RDCs, and ^13^Ca ^13^C RDCs, TALOS derived phi and psi restraints, hydrogen bonding restraints, ^3^J^HN-Ha^, 3J^Ha-Hβ^, and 3J^HN-Hβ^ J-couplings. Calculations were performed using NIH-XPLOR for minimizing the energies, each run generating 100 structures.

### SPR measurements

SPR datasets were generated using a BIAcore X100 instrument (GE Lifesciences, Piscataway, NJ) with biotinylated avi-tagged TBRI or biotinylated avi-tagged TBRII captured onto neutravidin-coated CM-5 sensor chips (GE Lifesciences, Piscataway, NJ) at a density of 50 – 150RU. Neutravidin coated sensor chips for capture of biotinylated avi-tag receptors were made by activating the surface of a CM-5 chip with EDC and NHS, followed by injection of neutravidin (Pierce, Rockford, IL) diluted into sodium acetate at pH 4.5 until the surface density reached 6000 – 15000 RU. Kinetic binding assays were performed by single injections of the analytes in 25 mM HEPES, pH 7.4, 50 mM NaCl, 0.005% surfactant P20 (Pierce, Rockford, IL) at 100 uL min^-1^. Regeneration of the surface was achieved by a 30 sec injection of 1 – 4 M guanidine hydrochloride. Baseline correction was performed by subtracting the response both from the reference surface with no immobilized ligand and 5 – 10 blank buffer injections. Kinetic analyses were performed by fitting the results to a simple 1:1 model using the program Scrubber (Biologic Software, Canberra, Australia).

### ITC measurements

ITC datasets were generated using a Microcal PEAQ-ITC instrument (Malvern Instruments, Westborough, MA). All experiments with TBRII were performed in ITC buffer, consisting of 25 mM sodium phosphate, 50 mM NaCl, pH 6.0 at 35 °C. Experiments with TBRI were performed in ITC buffer, consisting of 25 mM HEPES, 50mM NaCl, 0.05% NaN_3_, pH 7.5 at 25°C, with exception of the TBRI/TGF-B:TBRII titration which was performed at 30°C. A listing of the proteins included in the syringe and sample cell is provided in Tables 1-4.

Proteins included in the syringe and sample cell were dialyzed against ITC buffer and concentrated as necessary prior to being loaded into either the syringe or sample cell. For the TBRII experiments, fifteen 2.5 µL injections were performed with an injection duration of 5 s, a spacing of 150 s, and a reference power of 10. For the TBRI experiments, with exception of the TBRI/TGF-B:TBRII titration, nineteen 2.0 µL injections were performed with an injection duration of 4 s, a spacing of 150 s, and a reference power of 10. The TBRI/TGF-B:TBRII titration was performed with thirteen 3.0 µL injections were performed with an injection duration of 5 s, a spacing of 150 s, and a reference power of 10. Integration and data fitting were performed using the programs Nitpic^63^, Sedphat^64,65^, and GUSSI^66^. No more than 2 outlier data points were removed from any one ITC data set for analysis.

## Acknowledgements

The authors would like to thank Mike Delk for assistance with the NMR instrumentation. This research was supported by the NIH (GM58670) and the U.S. Department of Defense (DoD W81XWH-17-1-0429). Molecular graphics and analyses were performed with UCSF Chimera, which is developed by the Resource for Biocomputing, Visualization, and Informatics at the University of California, San Francisco and supported by NIGMS P41-GM103311.

## References

1 Pullan, R. L., Smith, J. L., Jasrasaria, R. & Brooker, S. J. Global numbers of infection and disease burden of soil transmitted helminth infections in 2010. Parasit Vectors 7, 37, doi:10.1186/1756-3305-7-37 (2014).

2 Hotez, P. J. & Molyneux, D. H. Tropical anemia: one of Africa’s great killers and a rationale for linking malaria and neglected tropical disease control to achieve a common goal. PLoS Negl Trop Dis 2, e270, doi:10.1371/journal.pntd.0000270 (2008).

3 Maizels, R. M., Smits, H. H. & McSorley, H. J. Modulation of Host Immunity by Helminths: The Expanding Repertoire of Parasite Effector Molecules. Immunity 49, 801–818, doi:10.1016/j.immuni.2018.10.016 (2018).

4 Ryan, S. M., Eichenberger, R. M., Ruscher, R., Giacomin, P. R. & Loukas, A. Harnessing helminth-driven immunoregulation in the search for novel therapeutic modalities. PLoS Pathog 16, e1008508, doi:10.1371/journal.ppat.1008508 (2020).

5 Wiedemann, M. & Voehringer, D. Immunomodulation and Immune Escape Strategies of Gastrointestinal Helminths and Schistosomes. Front Immunol 11, 572865, doi:10.3389/fimmu.2020.572865 (2020).

6 Maizels, R. M. & Smith, K. A. Regulatory T cells in infection. Adv Immunol 112, 73–136, doi:10.1016/B978-0-12-387827-4.00003-6 (2011).

7 Smith, K. A. et al.. Low-level regulatory T-cell activity is essential for functional type-2 effector immunity to expel gastrointestinal helminths. Mucosal Immunol 9, 428–443, doi:10.1038/mi.2015.73 (2016).

8 Chen, W. et al.. Conversion of peripheral CD4+CD25-naive T cells to CD4+CD25+ regulatory T cells by TGF-beta induction of transcription factor Foxp3. J Exp Med 198, 1875–1886, doi:10.1084/jem.20030152 (2003).

9 Sanjabi, S., Oh, S. A. & Li, M. O. Regulation of the Immune Response by TGF-beta: From Conception to Autoimmunity and Infection. Cold Spring Harb Perspect Biol 9, doi:10.1101/cshperspect.a022236 (2017).

10 Peng, Y., Laouar, Y., Li, M. O., Green, E. A. & Flavell, R. A. TGF-beta regulates in vivo expansion of Foxp3-expressing CD4+CD25+ regulatory T cells responsible for protection against diabetes. Proc Natl Acad Sci U S A 101, 4572–4577, doi:10.1073/pnas.0400810101 (2004).

11 Grainger, J. R. et al.. Helminth secretions induce de novo T cell Foxp3 expression and regulatory function through the TGF-beta pathway. J Exp Med 207, 2331–2341, doi:10.1084/jem.20101074 (2010).

12 Johnston, C. J. C. et al.. A structurally distinct TGF-beta mimic from an intestinal helminth parasite potently induces regulatory T cells. Nat Commun 8, 1741, doi:10.1038/s41467-017-01886-6 (2017).

13 Shull, M. M. et al.. Targeted disruption of the mouse transforming growth factor-beta 1 gene results in multifocal inflammatory disease. Nature 359, 693–699, doi:10.1038/359693a0 (1992).

14 Kaartinen, V. et al.. Abnormal lung development and cleft palate in mice lacking TGF-beta 3 indicates defects of epithelial-mesenchymal interaction. Nat Genet 11, 415–421, doi:10.1038/ng1295-415 (1995).

15 Sanford, L. P. et al.. TGFbeta2 knockout mice have multiple developmental defects that are non-overlapping with other TGFbeta knockout phenotypes. Development 124, 2659–2670 (1997).

16 Kelly, A., Houston, S. A., Sherwood, E., Casulli, J. & Travis, M. A. Regulation of Innate and Adaptive Immunity by TGFbeta. Adv Immunol 134, 137–233, doi:10.1016/bs.ai.2017.01.001 (2017).

17 Groppe, J. et al.. Cooperative assembly of TGF-beta superfamily signaling complexes is mediated by two disparate mechanisms and distinct modes of receptor binding. Mol Cell 29, 157–168, doi:10.1016/j.molcel.2007.11.039 (2008).

18 Wrana, J. L., Attisano, L., Wieser, R., Ventura, F. & Massague, J. Mechanism of activation of the TGF-beta receptor. Nature 370, 341–347, doi:10.1038/370341a0 (1994).

19 Wrana, J. L. et al.. TGF beta signals through a heteromeric protein kinase receptor complex. Cell 71, 1003–1014, doi:10.1016/0092-8674(92)90395-s (1992).

20 Ihara, S., Hirata, Y. & Koike, K. TGF-beta in inflammatory bowel disease: a key regulator of immune cells, epithelium, and the intestinal microbiota. J Gastroenterol 52, 777–787, doi:10.1007/s00535-017-1350-1 (2017).

21 Kim, K. K., Sheppard, D. & Chapman, H. A. TGF-beta1 Signaling and Tissue Fibrosis. Cold Spring Harb Perspect Biol 10, doi:10.1101/cshperspect.a022293 (2018).

22 Hu, H. H. et al.. New insights into TGF-beta/Smad signaling in tissue fibrosis. Chem Biol Interact 292, 76–83, doi:10.1016/j.cbi.2018.07.008 (2018).

23 Massague, J. TGFbeta in Cancer. Cell 134, 215–230, doi:10.1016/j.cell.2008.07.001 (2008).

24 Seoane, J. & Gomis, R. R. TGF-beta Family Signaling in Tumor Suppression and Cancer Progression. Cold Spring Harb Perspect Biol 9, doi:10.1101/cshperspect.a022277 (2017).

25 Mariathasan, S. et al. TGFbeta attenuates tumour response to PD-L1 blockade by contributing to exclusion of T cells. Nature 554, 544–548, doi:10.1038/nature25501 (2018).

26 Tauriello, D. V. F. et al. TGFbeta drives immune evasion in genetically reconstituted colon cancer metastasis. Nature 554, 538–543, doi:10.1038/nature25492 (2018).

27 Li, S. et al. Cancer immunotherapy via targeted TGF-beta signalling blockade in TH cells. Nature 587, 121–125, doi:10.1038/s41586-020-2850-3 (2020).

28 Kirkitadze, M. D. & Barlow, P. N. Structure and flexibility of the multiple domain proteins that regulate complement activation. Immunol Rev 180, 146–161, doi:10.1034/j.1600-065x.2001.1800113.x (2001).

29 Smyth, D. J. et al. TGF-beta mimic proteins form an extended gene family in the murine parasite Heligmosomoides polygyrus. Int J Parasitol 48, 379–385, doi:10.1016/j.ijpara.2017.12.004 (2018).

30 Hewitson, J. P. et al. Secretion of protective antigens by tissue-stage nematode larvae revealed by proteomic analysis and vaccination-induced sterile immunity. PLoS Pathog 9, e1003492, doi:10.1371/journal.ppat.1003492 (2013).

31 Osbourn, M. et al. HpARI Protein Secreted by a Helminth Parasite Suppresses Interleukin-33. Immunity 47, 739–751 e735, doi:10.1016/j.immuni.2017.09.015 (2017).

32 Chauche, C. et al. A Truncated Form of HpARI Stabilizes IL-33, Amplifying Responses to the Cytokine. Front Immunol 11, 1363, doi:10.3389/fimmu.2020.01363 (2020).

33 Vacca, F. et al. A helminth-derived suppressor of ST2 blocks allergic responses. Elife 9, doi:10.7554/eLife.54017 (2020).

34 Zuniga, J. E. et al. Assembly of TbetaRI:TbetaRII:TGFbeta ternary complex in vitro with receptor extracellular domains is cooperative and isoform-dependent. J Mol Biol 354, 1052–1068, doi:10.1016/j.jmb.2005.10.014 (2005).

35 Inman, G. J. et al. SB-431542 is a potent and specific inhibitor of transforming growth factor-beta superfamily type I activin receptor-like kinase (ALK) receptors ALK4, ALK5, and ALK7. Mol Pharmacol 62, 65–74, doi:10.1124/mol.62.1.65 (2002).

36 Gaboriaud, C., Rossi, V., Bally, I., Arlaud, G. J. & Fontecilla-Camps, J. C. Crystal structure of the catalytic domain of human complement c1s: a serine protease with a handle. EMBO J 19, 1755–1765, doi:10.1093/emboj/19.8.1755 (2000).

37 Reid, K. B. & Day, A. J. Structure-function relationships of the complement components. Immunol Today 10, 177–180, doi:10.1016/0167-5699(89)90317-4 (1989).

38 Gerhard Wider, D. N., Kurt Wuthrich. Studies of slow conformational equilibria in macromolecules by exchange of heteronuclear longitudinal 2-spin-order in a 2D difference correlation experiment. Journal of Biomolecular NMR 1, 5, doi:10.1007/BF01874572 (1991).

39 Kim, S. K. et al. An engineered transforming growth factor beta (TGF-beta) monomer that functions as a dominant negative to block TGF-beta signaling. J Biol Chem 292, 7173–7188, doi:10.1074/jbc.M116.768754 (2017).

40 Hinck, A. P. et al. Sequential resonance assignments of the extracellular ligand binding domain of the human TGF-beta type II receptor. J Biomol NMR 18, 369–370 (2000).

41 Ilangovan, U., Deep, S., Hinck, C. S. & Hinck, A. P. Sequential resonance assignments of the extracellular domain of the human TGFbeta type II receptor in complex with monomeric TGFbeta3. J Biomol NMR 29, 103–104, doi:10.1023/B:JNMR.0000019468.50957.42 (2004).

42 Eghbalnia, H. R., Wang, L., Bahrami, A., Assadi, A. & Markley, J. L. Protein energetic conformational analysis from NMR chemical shifts (PECAN) and its use in determining secondary structural elements. J Biomol NMR 32, 71–81, doi:10.1007/s10858-005-5705-1 (2005).

43 Holm, L. Using Dali for Protein Structure Comparison. Methods Mol Biol 2112, 29–42, doi:10.1007/978-1-0716-0270-6_3 (2020).

44 Holm, L. DALI and the persistence of protein shape. Protein Sci 29, 128–140, doi:10.1002/pro.3749 (2020).

45 Radaev, S. et al. Ternary complex of transforming growth factor-beta1 reveals isoform-specific ligand recognition and receptor recruitment in the superfamily. J Biol Chem 285, 14806–14814, doi:10.1074/jbc.M109.079921 (2010).

46 Hart, P. J. et al. Crystal structure of the human TbetaR2 ectodomain--TGF-beta3 complex. Nat Struct Biol 9, 203–208, doi:10.1038/nsb766 (2002).

47 McNeilly, T. N. & Nisbet, A. J. Immune modulation by helminth parasites of ruminants: implications for vaccine development and host immune competence. Parasite 21, 51, doi:10.1051/parasite/2014051 (2014).

48 Geiger, S. M. et al. Stage-specific immune responses in human Necator americanus infection. Parasite Immunol 29, 347–358, doi:10.1111/j.1365-3024.2007.00950.x (2007).

49 Bower, M. A., Constant, S. L. & Mendez, S. Necator americanus: the Na-ASP-2 protein secreted by the infective larvae induces neutrophil recruitment in vivo and in vitro. Exp Parasitol 118, 569–575, doi:10.1016/j.exppara.2007.11.014 (2008).

50 Teixeira-Carvalho, A. et al. Binding of excreted and/or secreted products of adult hookworms to human NK cells in Necator americanus-infected individuals from Brazil. Infect Immun 76, 5810–5816, doi:10.1128/IAI.00419-08 (2008).

51 Kelley, L. A., Mezulis, S., Yates, C. M., Wass, M. N. & Sternberg, M. J. The Phyre2 web portal for protein modeling, prediction and analysis. Nat Protoc 10, 845–858, doi:10.1038/nprot.2015.053 (2015).

52 Zuniga, J. E. et al. The TbetaR-I pre-helix extension is structurally ordered in the unbound form and its flanking prolines are essential for binding. J Mol Biol 412, 601–618, doi:10.1016/j.jmb.2011.07.046 (2011).

53 Walker, R. G. et al. Structural basis for potency differences between GDF8 and GDF11. BMC Biol 15, 19, doi:10.1186/s12915-017-0350-1 (2017).

54 Kay, L. E., Keifer, P., Saarinen, T. Journal of the American Chemical Society, 10663–10665 (1992).

55 Grzesiek, S., Bax, A,. The importance of not saturating water in protein NMR. Application to sensitivity enhancement and NOE measurements. Journal of the American Chemical Society 115, 12593–12594, doi:10.1021/ja00079a052 (1993).

56 Pitto, M., Saudek, V., Sklenar, V. Gradient-tailored excitation for single-quantum NMR spectroscopy of aqueous solutions. Journal of Biomolecular NMR 2, 661–665 (1992).

57 Delaglio, F. et al. NMRPipe: a multidimensional spectral processing system based on UNIX pipes. J Biomol NMR 6, 277–293 (1995).

58 Vranken, W. F. et al. The CCPN data model for NMR spectroscopy: development of a software pipeline. Proteins 59, 687–696, doi:10.1002/prot.20449 (2005).

59 Bahrami, A., Assadi, A. H., Markley, J. L. & Eghbalnia, H. R. Probabilistic interaction network of evidence algorithm and its application to complete labeling of peak lists from protein NMR spectroscopy. PLoS Comput Biol 5, e1000307, doi:10.1371/journal.pcbi.1000307 (2009).

60 Montelione, G. T. et al. Sequence-specific 1H-NMR assignments and identification of two small antiparallel beta-sheets in the solution structure of recombinant human transforming growth factor alpha. Proc Natl Acad Sci U S A 86, 1519–1523, doi:10.1073/pnas.86.5.1519 (1989).

61 Ottiger, M., Delaglio, F. & Bax, A. Measurement of J and dipolar couplings from simplified two-dimensional NMR spectra. J Magn Reson 131, 373–378, doi:10.1006/jmre.1998.1361 (1998).

62 Schwieters, C. D., Kuszewski, J. J., Tjandra, N. & Clore, G. M. The Xplor-NIH NMR molecular structure determination package. J Magn Reson 160, 65–73, doi:10.1016/s1090-7807(02)00014-9 (2003).

63 Keller, S. et al. High-precision isothermal titration calorimetry with automated peak-shape analysis. Anal Chem 84, 5066–5073, doi:10.1021/ac3007522 (2012).

64 Brautigam, C. A., Zhao, H., Vargas, C., Keller, S. & Schuck, P. Integration and global analysis of isothermal titration calorimetry data for studying macromolecular interactions. Nat Protoc 11, 882–894, doi:10.1038/nprot.2016.044 (2016).

65 Zhao, H., Piszczek, G. & Schuck, P. SEDPHAT--a platform for global ITC analysis and global multi-method analysis of molecular interactions. Methods 76, 137–148, doi:10.1016/j.ymeth.2014.11.012 (2015).

66 Brautigam, C. A. Calculations and Publication-Quality Illustrations for Analytical Ultracentrifugation Data. Methods Enzymol 562, 109–133, doi:10.1016/bs.mie.2015.05.001 (2015).

