## Supplementary Information for "Structure-based mapping of the TβRI and TβRII receptor binding sites of the parasitic TGF-β mimic, Hp-TGM"

**Table S1. *H. polygyrus* constructs used in this study**

| Construct | Residue range and features* | Sequence |
|---|---|---|
| TGM-D1 | Residues 16-95 of *H. polygyrus* TGF-β Mimic (NCBI ATO59092.1)<br><br>Expressed as a Thioredoxin-fusion<br><br>GT from KpnI cut site | MSDKIIHLTDDSFDTDVLKADGAILVDFWA EWCGPCKMIAPILDEIADEYQGKLTVAKLN IDQNPGTAPKYGIRGIPTLLLFKNGEVAATK VGALSKGQLKEFLDANLAGSGSGHMHHHH HHSSGLVPRGS GTGSGSGSDDSGCMPFSDE AATYKYVAKGPKNIEIPAQIDNSGMYPDYT HVKRFCKGLHGEDTTGWFVGICLASQWYY YEGVQECDDR |
| TGM-D2 | Residues 96-176 of *H. polygyrus* TGF-β Mimic (NCBI ATO59092.1)<br><br>Expressed as a Thioredoxin-fusion<br><br>GT from KpnI cut site | MSDKIIHLTDDSFDTDVLKADGAILVDFWA EWCGPCKMIAPILDEIADEYQGKLTVAKLN IDQNPGTAPKYGIRGIPTLLLFKNGEVAATK VGALSKGQLKEFLDANLAGSGSGHMHHHH HHSSGLVPRGS GTRCSPLPTNDTVSFEYLK ATVNPGIIFNITVHPDASGKYPELTYIKRICK NFPTDSNVQGHIIGMCYNAEWQFSSTPTCP AS |
| TGM-D1D2 | Residues 16-176 of *H. Polygyrus* TGF-β Mimic (NCBI ATO59092.1)<br><br>Expressed as a Thioredoxin-fusion<br><br>GT from KpnI cut site | MSDKIIHLTDDSFDTDVLKADGAILVDFWA EWCGPCKMIAPILDEIADEYQGKLTVAKLN IDQNPGTAPKYGIRGIPTLLLFKNGEVAATK VGALSKGQLKEFLDANLAGSGSGHMHHHH HHSSGLVPRGS GTGSGSGSDDSGCMPFSDE AATYKYVAKGPKNIEIPAQIDNSGMYPDYT HVKRFCKGLHGEDTTGWFVGICLASQWYY YEGVQECDDRRCSPLPTNDTVSFEYLKATV NPGIIFNITVHPDASGKYPELTYIKRICKNFP TDSNVQGHIIGMCYNAEWQFSSTPTCPAS |
| TGM-D3 | Residues 177-262 of *H. Polygyrus* TGF-β Mimic (NCBI ATO59092.1)<br><br>Expressed as a Thioredoxin-fusion<br><br>GT from KpnI cut site | MSDKIIHLTDDSFDTDVLKADGAILVDFWA EWCGPCKMIAPILDEIADEYQGKLTVAKLN IDQNPGTAPKYGIRGIPTLLLFKNGEVAATK VGALSKGQLKEFLDANLAGSGSGHMHHHH HHSSGLVPRGS GTGCPPLPDDGIVFYEYYG YAGDRHTVGPVVTKDSSGNYPSPTHARRR CRALSQEADPGEFVAICYKSGTTGESHWEY YKNIGKCPDP |
| TGM-FL | Residues 16-422 of *H. Polygyrus* TGF-β Mimic (NCBI ATO59092.1)<br><br>Signal Peptide<br>pSECTag + terminal His tag | METDTLLLWVLLLWVPGSTGDAAQPARRA DDSGCMPFSDEAATYKYVAKGPKNIEIPAQI DNSGMYPDYTHVKRFCKGLHGEDTTGWFV GICLASQWYYYEGVQECDDRRCSPLPTNDTV SFEYLKATVNPGIIFNITVHPDASGKYPELTYI KRICKNFPTDSNVQGHIIGMCYNAEWQFSST PTCPASGCPPLPDDGIVFYEYYGYAGDRHTV GPVVTKDSSGNYPSPTHARRRCRALSQEAD PGEFVAICYKSGTTGESHWEYYKNIGKCPD |

| | | |
|---|---|---|
| | | PRCKPLEANESVHYEYFTMTNETDKKKGPP AKVGKSGKYPEHTCVKKVCSKWPYTCSTG GPIFGECIGATWNFTALMECINARGCSSDDL FDKLGFEKVIVRKGEGSDSYKDDFARFYAT GSKVIAECGGKTVRLECSNGEWHEPGTKTV HRCTKDGIRTLGPEQKLISEEDLNSAVDHHH HHH |

**Table S2. Receptor and Growth Factor constructs used in this study**

| Construct | Residue range and features* | Sequence |
|---|---|---|
| TβRI | Residues 25-125 of the human TGF-β type I receptor (NCBI NP_004603.1)<br><br>His-Tag<br>Thrombin-Cleavage Site<br>HM tag | MGSSHHHHHHSSGLVPRGSHMAALLPG ATALQCFCHLCTKDNFTCVTDGLCFVSV TETTDKVIHNSSCIAEIDLIPRDRPFVCAP SSKTGSVTTTYCCNQDHCNKIELPTTVK SSPGLGPVE |
| TβRII | Residues 38-153 of the human TGF-β type II receptor (NCBI AHI94913) | MVTDNNGAVKFPQLCKFCDVRFSTCDQ KSCMSNCSITSICEKPQEVCVAVWRKNE NITLETVCHDPKLPYHDFILEDAASPKCI MKEKKKPGETFFMCSCSSDECNDNIIFSE EY |
| mmTGF-β2 7M2R | Residues 303-352 and 377-414 of mouse TGF-β2 (NCBI NP_0033393) connected by an engineered loop<br><br>C379R substitution render the protein monomeric; K327R, R328K, V381R, L391V, I394V, K396R, T397K, and I400V substitutions enable high affinity TβRII binding and high solubility | ALDAAYCFRNVQDNCCLRPLYIDFRKD LGWKWIHEPKGYNANFCAGACPYRAS KSPRCRSQDLEPLTIVYYVGRKPKVEQ LSNMIVKSCKCS |

*All residue numbering begins with the N-terminal methionine of the naturally occurring signal peptide

**Table S3: SPR Binding constants for TβRII and TGM domains**

| Surface | Analyte | Dissociation constant | | | |
|---|---|---|---|---|---|
| | | $k_{on}$ ($M^{-1}$ $s^{-1}$) | $k_{off}$ ($s^{-1}$) | $K_d$ (μM) | $R_{max}$ (RU) |
| TβRII | TGM D1 | ND[a] | ND[a] | ND[a] | ND[a] |
| TβRII | TGM D2 | ND[a] | ND[a] | ND[a] | ND[a] |
| TβRII | TGM D1D2 | ND[a] | ND[a] | ND[a] | ND[a] |
| TβRII | TGM D3 | $(6 \pm 1) \times 10^5$ | $0.6 \pm 0.1$ | $0.91 \pm 0.02$ | $33.0 \pm 0.4$ |
| TβRII | TGM FL | $(2 \pm 6) \times 10^7$ | $(1 \pm 4) \times 10^{-1}$ | $0.61 \pm 0.01$ | $215 \pm 2$ |

[a]Not determined due to weak signal

**Table S4. TGM:TβRI binding as assessed by SPR**

| Surface | Analyte | Fitted Parameters | | | |
|---|---|---|---|---|---|
| | | $k_{on}$ ($M^{-1}$ $s^{-1}$) | $k_{off}$ ($s^{-1}$) | $K_d$ (nM) | $R_{max}$ (RU) |
| TβRI | TGM-D1 | ND[a] | ND[a] | ND[a] | ND[a] |
| TβRI | TGM-D2 | $(2.95 \pm 0.04) \times 10^5$ | $(9.11 \pm 0.07) \times 10^{-2}$ | $309 \pm 4$ | $89.6 \pm 0.7$ |
| TβRI | TGM-D3 | ND[a] | ND[a] | ND[a] | ND[a] |
| TβRI | TGM-D1D2 | $(6.72 \pm 0.01) \times 10^4$ | $(1.62 \pm 0.01) \times 10^{-3}$ | $24.1 \pm 0.1$ | $429 \pm 1$ |
| TβRI[b] | TGM-FL | $(5.93 \pm 0.02) \times 10^4$ | $(7.8 \pm 0.2) \times 10^{-4}$ | $13.1 \pm 0.4$ | $193 \pm 1$ |

[a]Not determined due to weak signal

[b]Low density chip

**Figure S1**. **$^1$H-$^{15}$N HSQC spectra of TGM-D3 and TGM-D2**. A. $^1$H-$^{15}$N HSQC spectrum of $^{15}$N TGM-D3. B-C. $^1$H-$^{15}$N HSQC spectrum of $^{15}$N TGM-D2 (B). Blue boxes mark doubled peaks in dynamic equilibrium with one another as identified by a ZZ-exchange HSQC experiment (expansion of ZZ-exchange HSQC spectrum with a mixing time of 250 ms is shown as an inset for two pairs of peaks). Peak expansion corresponding to ZZ-exchange HSQC experiment as a function of the mixing time is shown for the pair of peaks at $^1$H 10.3 ppm/$^{15}$N 124 ppm (C). All spectra recorded in 25 mM sodium phosphate, 50 mM sodium chloride, 5% $^2H_2O$ pH 6.0, 310 K.

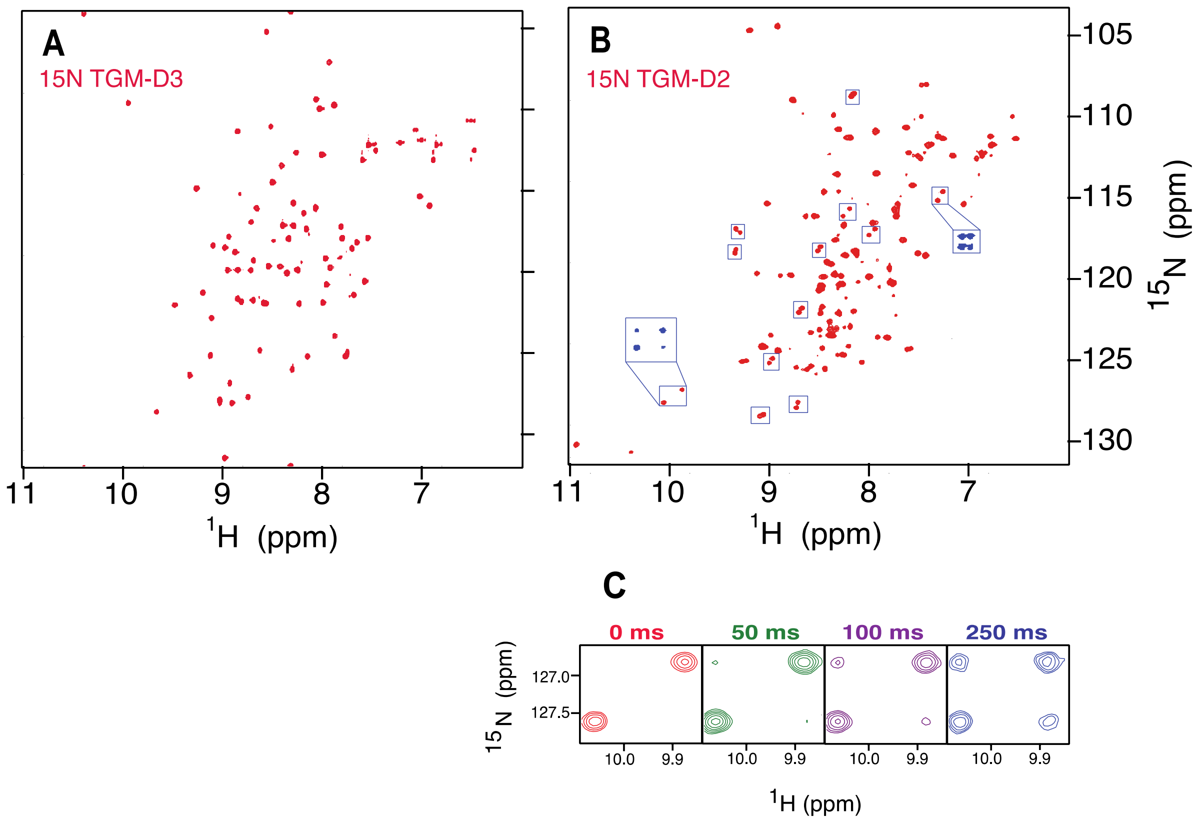

**Figure S2**. **$^{1}$H-$^{15}$N HSQC spectra of TGM-D1**. A. $^{1}$H-$^{15}$N HSQC spectrum of 100 μM $^{15}$N TGM-D1 in 25 mM sodium phosphate, 250 mM sodium chloride, 5% $^{2}H_2O$ pH 6.0, 310 K. B. $^{1}$H-$^{15}$N HSQC spectrum of 200 μM $^{15}$N TGM-D1 with 10 mM CHAPS added. C-D. $^{1}$H-$^{15}$N HSQC spectra of 20 μM $^{15}$N TGM-D1 in the same buffer as in Panel A (C) or with 10 mM CHAPS added (D).

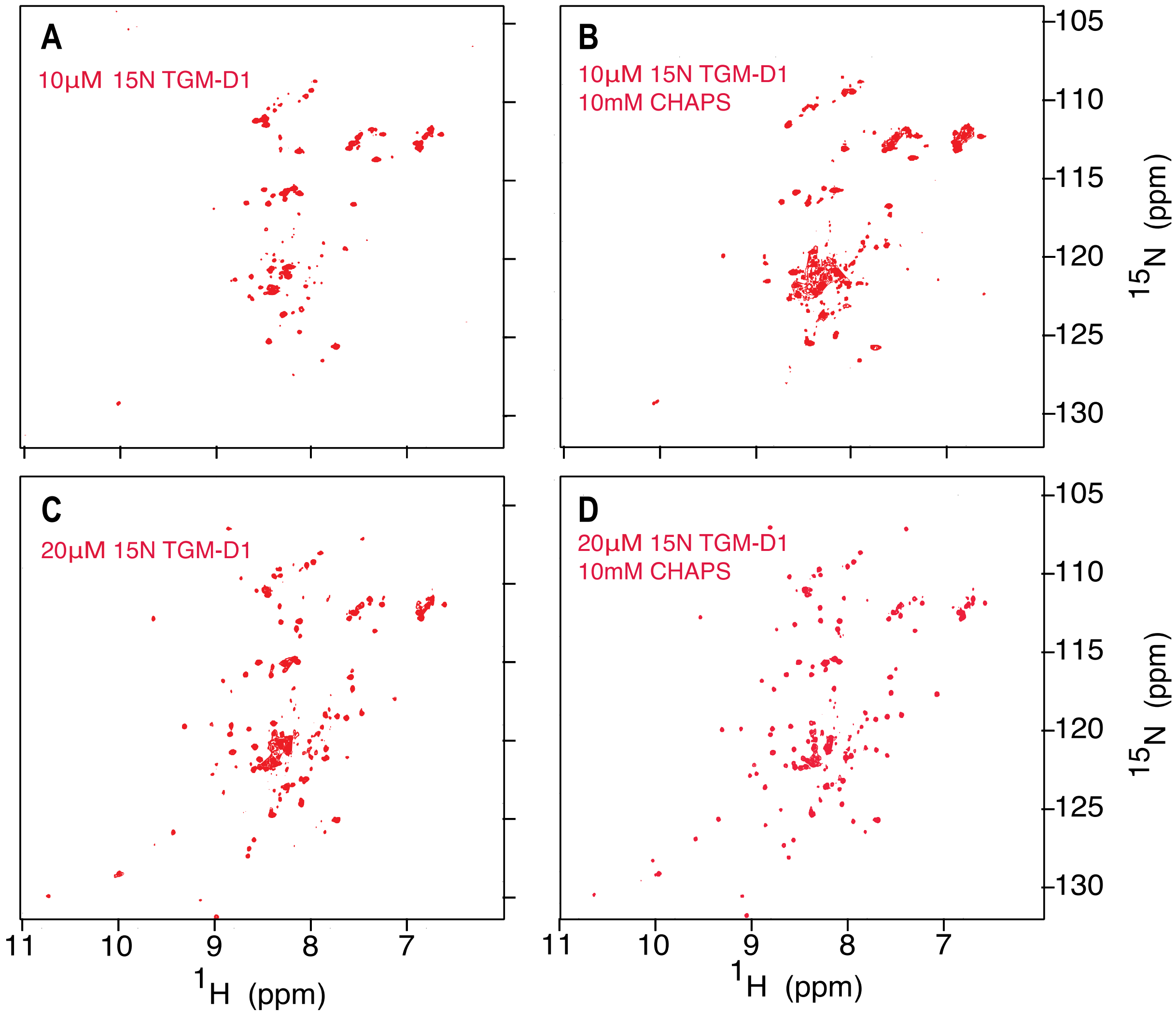

**Figure S3. Binding of TβRI TGM-D1 and TGM-D3 by and of TβRII TGM-D1 and TGM-D2.** A-B. $^{1}H$-$^{15}N$ HSQC spectra 0.04 mM $^{15}N$ TGM-D1 (A) or 0.1 mM $^{15}N$ TGM-D3 (B) alone (red) overlaid with the spectrum of the same sample but with an excess of unlabeled TβRII (1.2 molar equivalents) added (blue). C-D. $^{1}H$-$^{15}N$ HSQC spectra 0.04 mM $^{15}N$ TGM-D1 (C) or 0.2 mM $^{15}N$ TGM-D3 (D) alone (red) overlaid with the spectrum of the same sample but with an excess of unlabeled TβRI (1.2 molar equivalents) added (blue). Spectra were recorded in 25 mM sodium phosphate, 50 mM sodium chloride, 5% $^{2}H_2O$ pH 6.0, 310 K in either the presence of (A,C) or absence of 10 mM CHAPS (B, D).

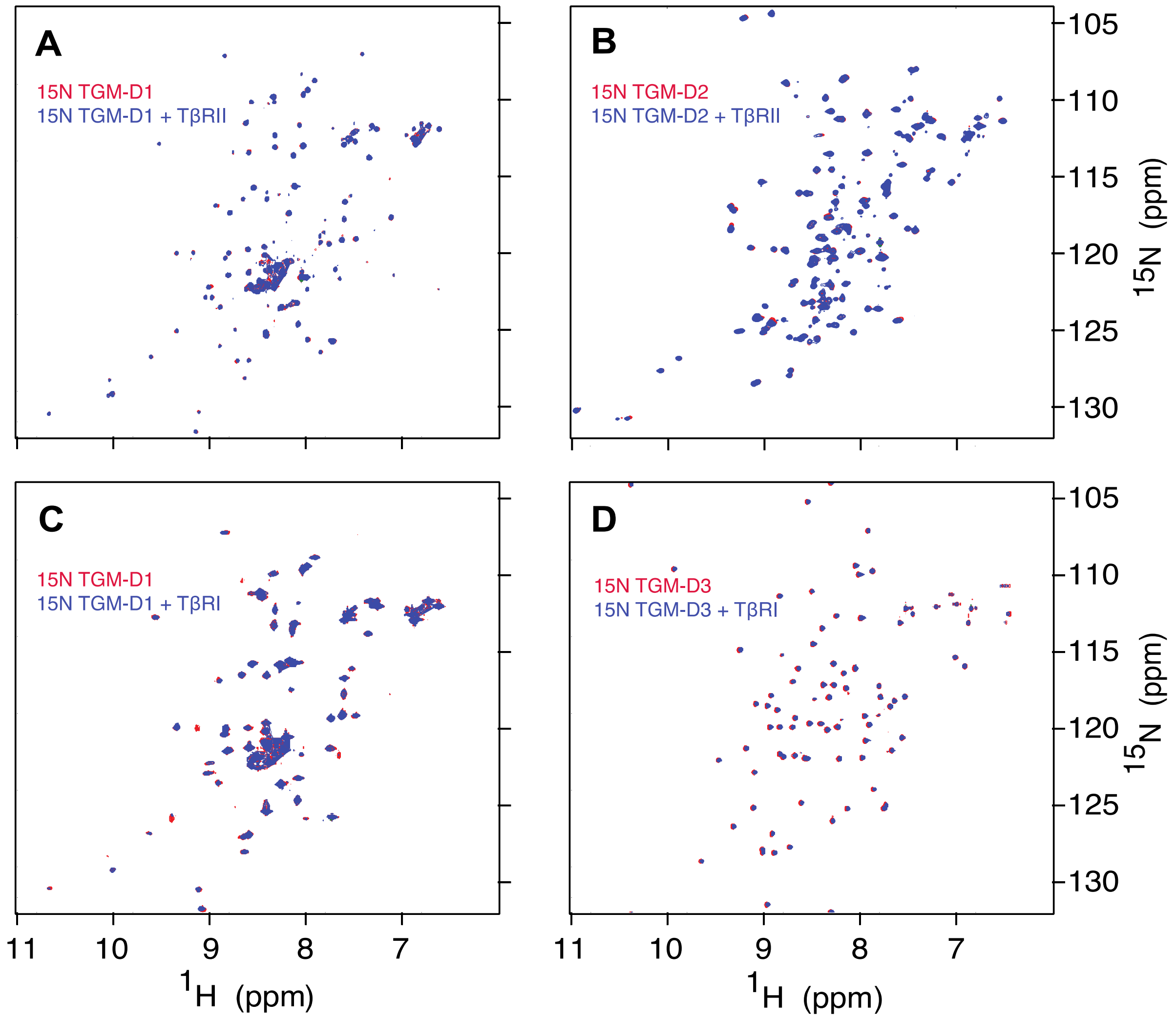

**Figure S4. NMR titrations of TβRI by TGM-D2 and associated SEC. A-D.** $^{1}H$-$^{15}N$ HSQC regions for correlating to expansions of the boxed regions in Figure 1A with all titration points: $^{15}N$ TGM-D2 alone (A, red) with 40µL TβRI stock (B, green), 160µL TβRI stock (C, purple), or 280 µL TβRI stock added (D, blue). **E-H.** Corresponding SEC (Superdex 75 10/300) chromatograms correlating to the titration points: $^{15}N$ TGM-D2 alone (E, red) with 40µL TβRI stock (F, green), 160µL TβRI stock (G, purple), or 280 µL TβRI stock added (H, blue). The dashed line connecting the panels of E-H is a chromatogram run with 150µL TβRI stock alone. All spectra were recorded in 25 mM sodium phosphate, 50 mM sodium chloride, 5% $^{2}H_2O$ pH 7.0, 310 K. **I.** SDS-PAGE gel of peaks marked in panel G with marker displayed as the left-most lane. **J.** SEC-MALS (Superdex 75 10/300) chromatogram correlating TβRI:TGM-D2 complex formation to molecular mass of displayed peaks.

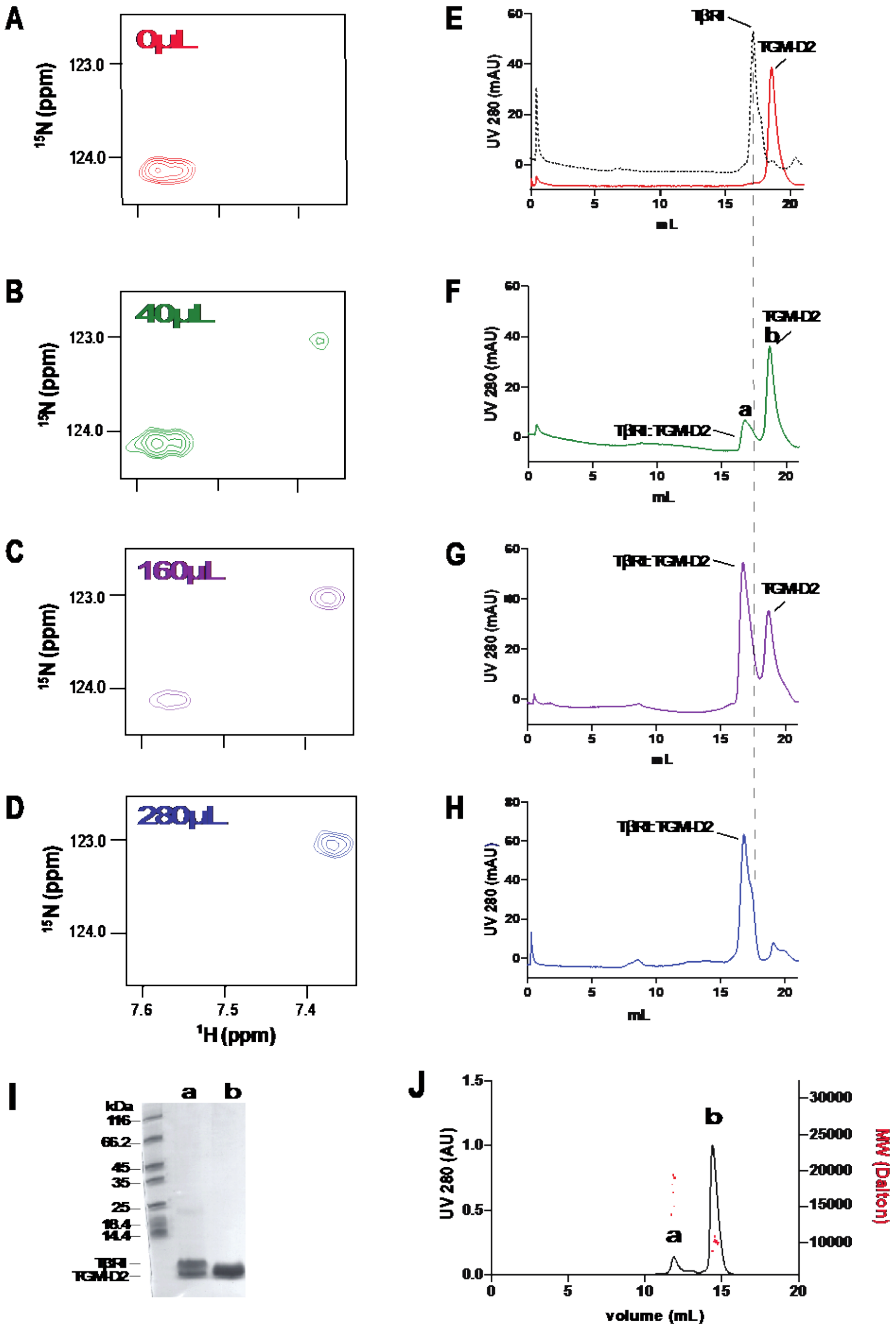

**Figure S5. Binding of TGM-D2 by TβRI.** A-B. $^{1}H$-$^{15}N$ HSQC spectra of TGM-D2 alone (A) or with an excess of unlabeled TβRI (B). All spectra recorded in 25 mM sodium phosphate, 50 mM sodium chloride, 5% $^{2}H_2O$ pH 6.0, 310 K. The boxed regions on the spectra mark peaks in conformational exchange (A) or resolved doubled peaks (B).

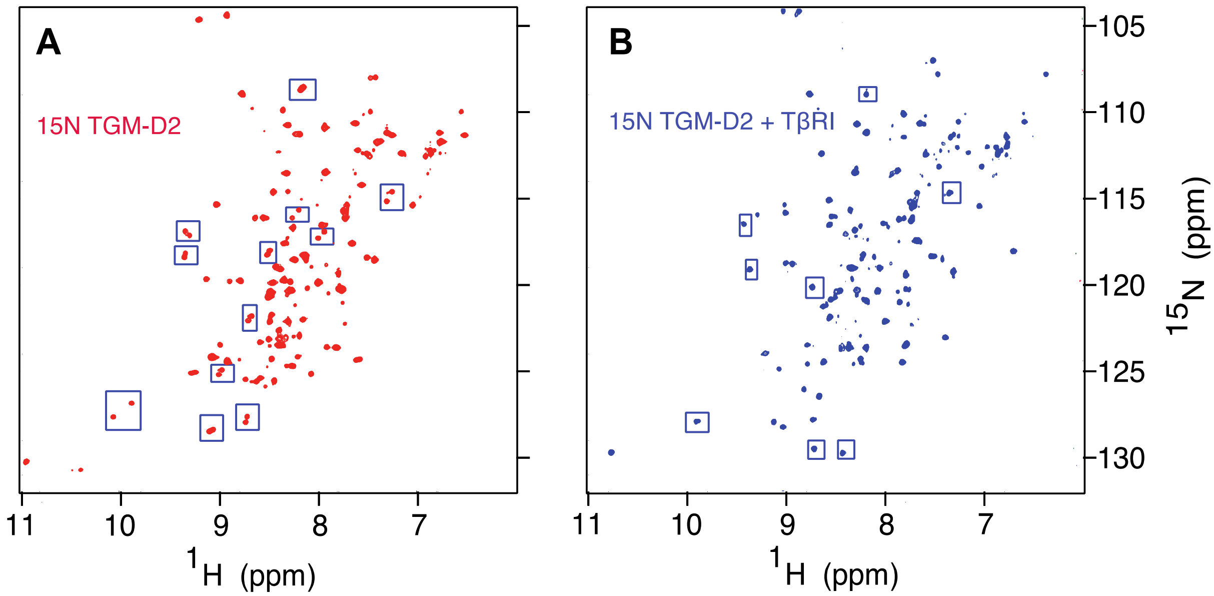

**Figure S6. Binding of TβRII by TGM-D3 and of TβRI by TGM-D2, respectively.** A-B. $^{15}$N HSQC spectra of 0.03 mM $^{15}$N TβRII alone (red) overlaid with the spectrum of the same sample, but with an excess of unlabeled TGM-D3 (blue) (A) or $^{1}$H-$^{15}$N HSQC spectra of 150 μL $^{15}$N TβRI stock alone (red) overlaid with the spectrum of the same sample, but with an excess of unlabeled TGM-D2 (blue) (B). Spectra recorded in 25 mM sodium phosphate, 50 mM sodium chloride, 5% $^{2}H_2O$ pH 6.0, 310 K (A) or 300K (B). Expansion of the boxed region of the spectra shown in panels A and B with immediate titration points is shown below in panels C and D, respectively, where numbers either indicate molar equivalents of TGM-D3 added (C) or volume of 797μM TGM-D2 added (μL) (D).

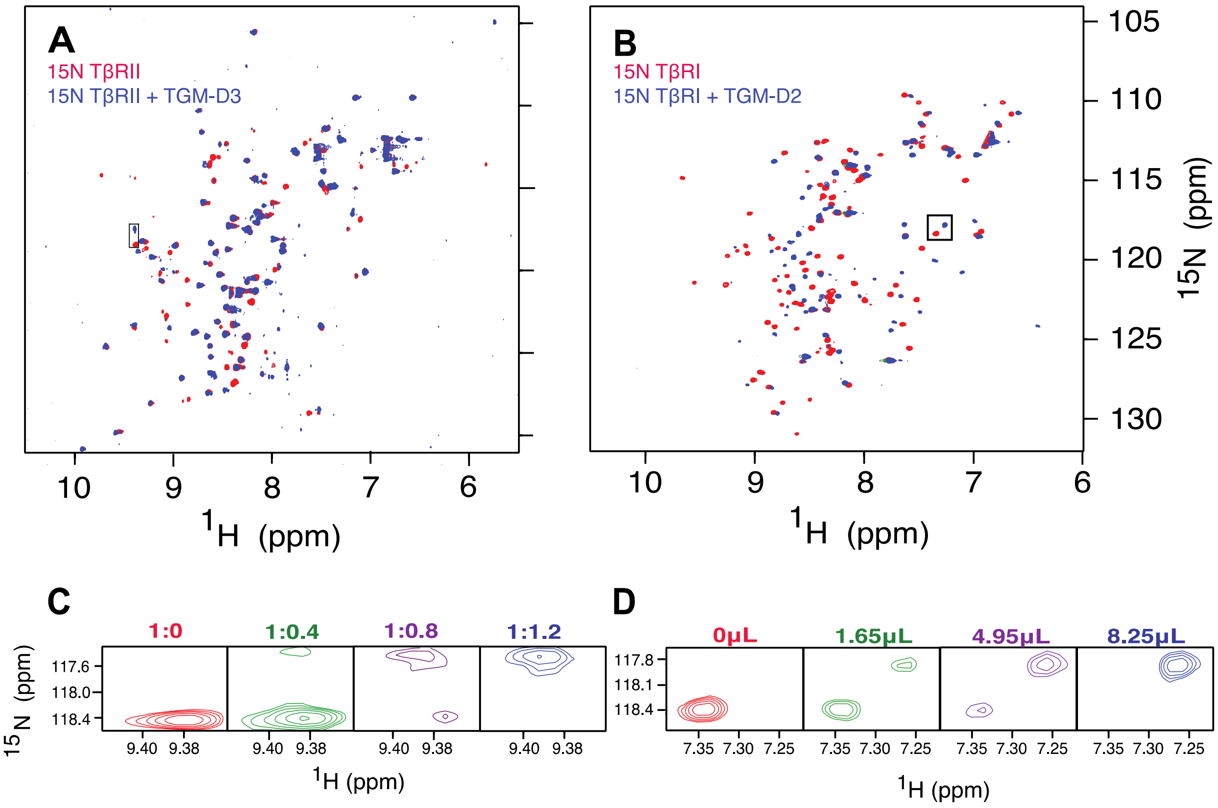

**Figure S7**. **Binding of TGM-D1, and TGM-D2 by TβRII.** A-B. $^{1}H$-$^{15}N$ HSQC spectra 0.03 mM $^{15}N$ TβRII alone (red) overlaid with the spectrum of the same sample but with an excess of unlabeled TGM-D1 (A) (1.5 molar equivalents) or TGM-D2 (B) (1.5 molar equivalents) added (blue). Spectra were recorded in 25 mM sodium phosphate, 50 mM sodium chloride, 5% $^{2}H_2O$ pH 6.0, 310 K.

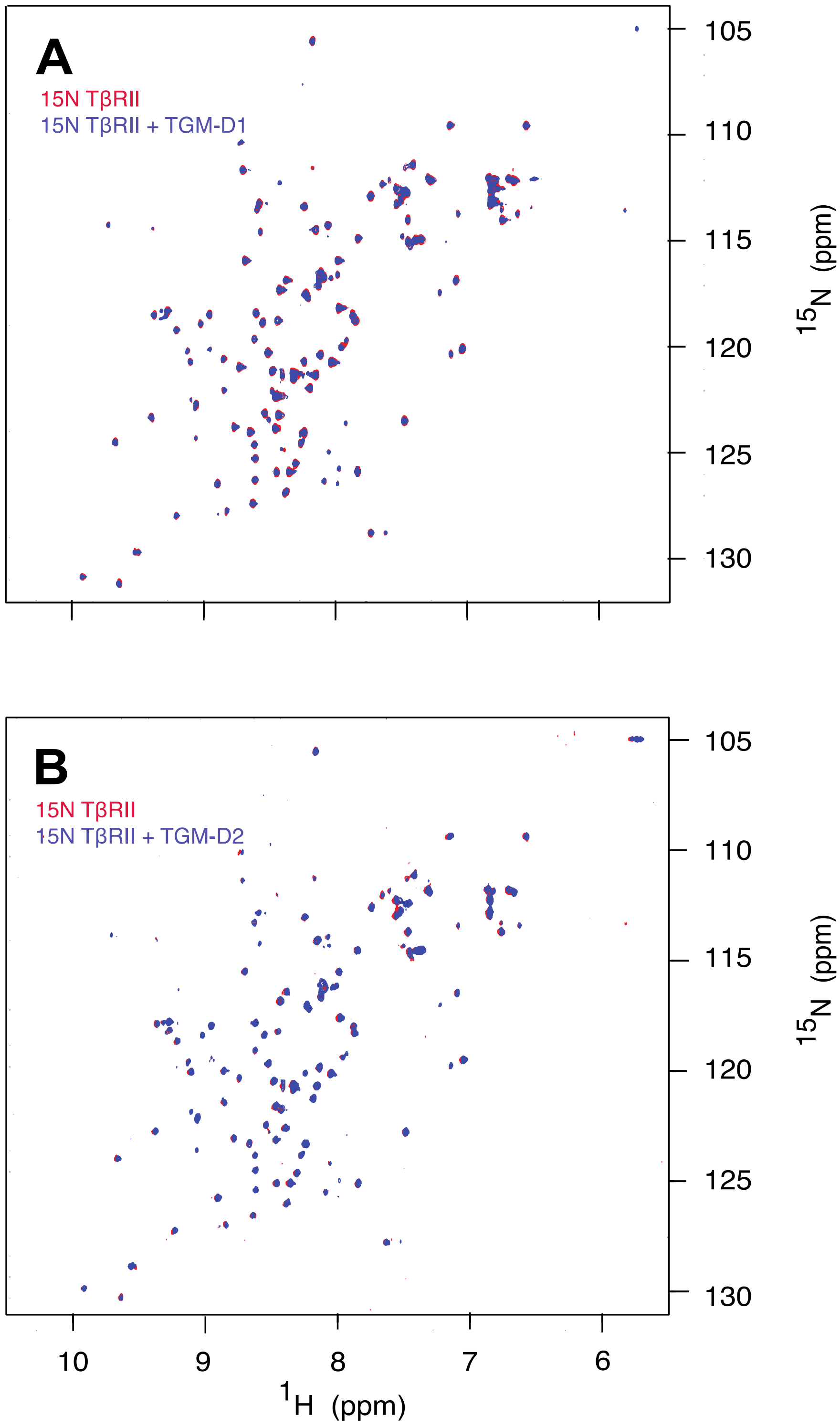

**Figure S8**. **Binding of TGM-D3 and TGM-D1 by TβRI.** A-B. $^{1}H$-$^{15}N$ HSQC spectra 0.02 mM $^{15}N$ TβRI alone (red) overlaid with the spectrum of the same sample but with an excess of unlabeled TGM-D3 (A) (3.0 molar equivalents) or TGM-D1 (B) (3.0 molar equivalents) added (blue). Spectra were recorded in 25 mM sodium phosphate, 50 mM sodium chloride, 5% $^{2}H_2O$ pH 6.6, 300 K.

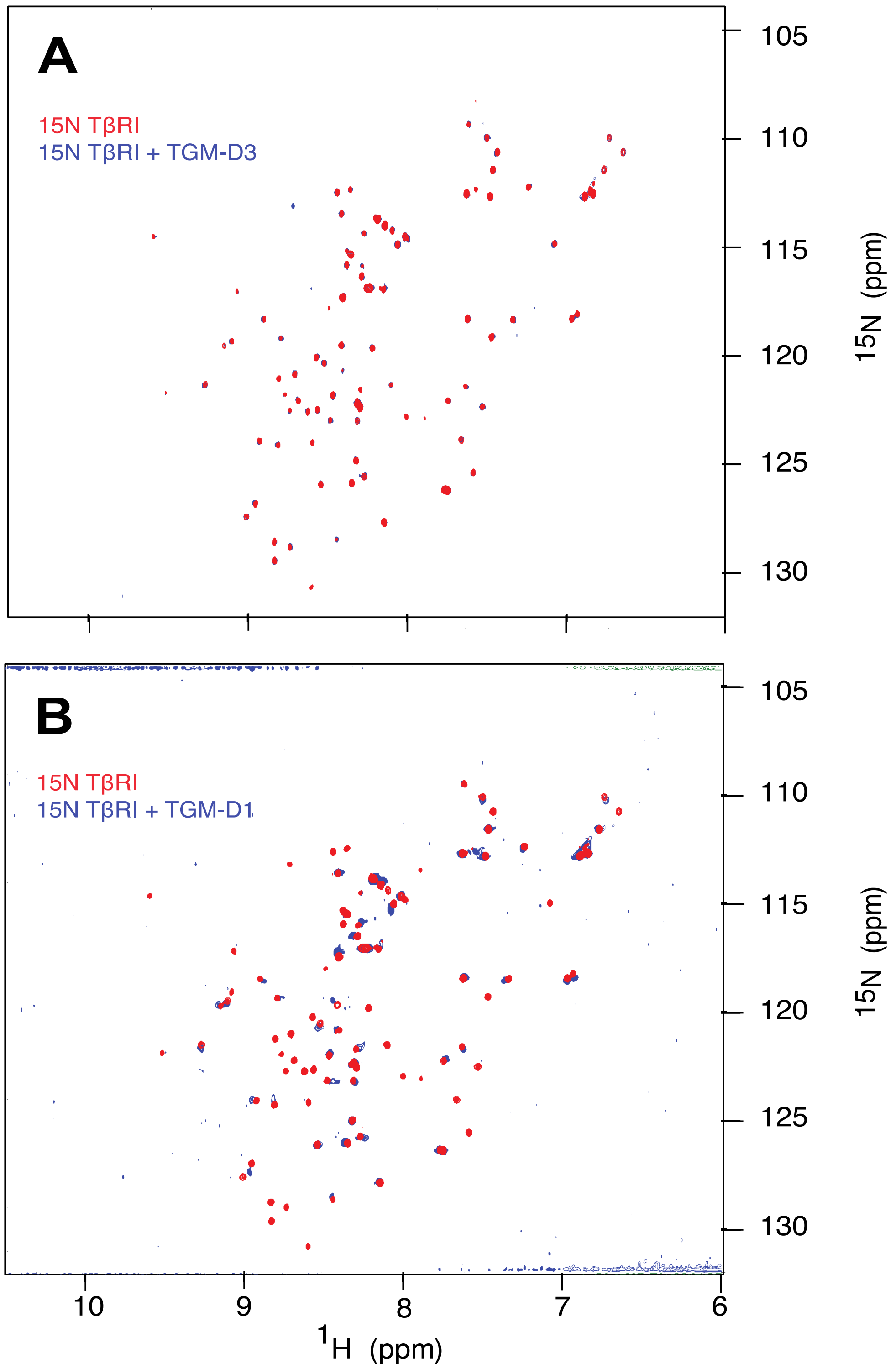

**Figure S9. $^{1}H$-$^{15}N$ HSQC Assignments of TβRII alone and bound to TGM-D3.** A. $^{1}H$-$^{15}N$ HSQC spectra of TβRII alone with peaks assigned. B. $^{1}H$-$^{15}N$ HSQC spectra of TβRII bound to TGM-D3 with peaks assigned. Dashed horizontal lines indicate Nε/Hε pairs. Spectra recorded in 25 mM sodium phosphate, 50 mM sodium chloride, 5% $^{2}H_2O$ pH 6.0, 310K

**A**

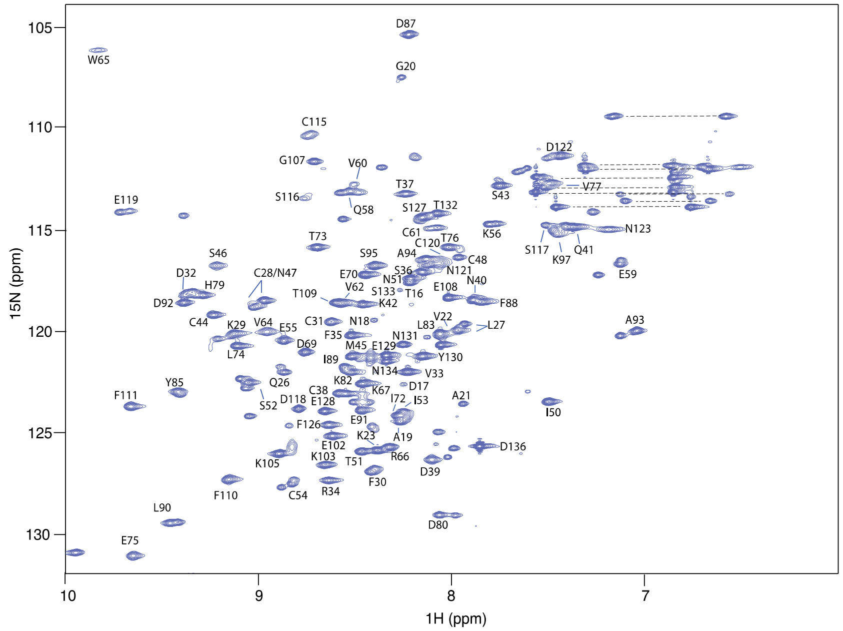


**B**

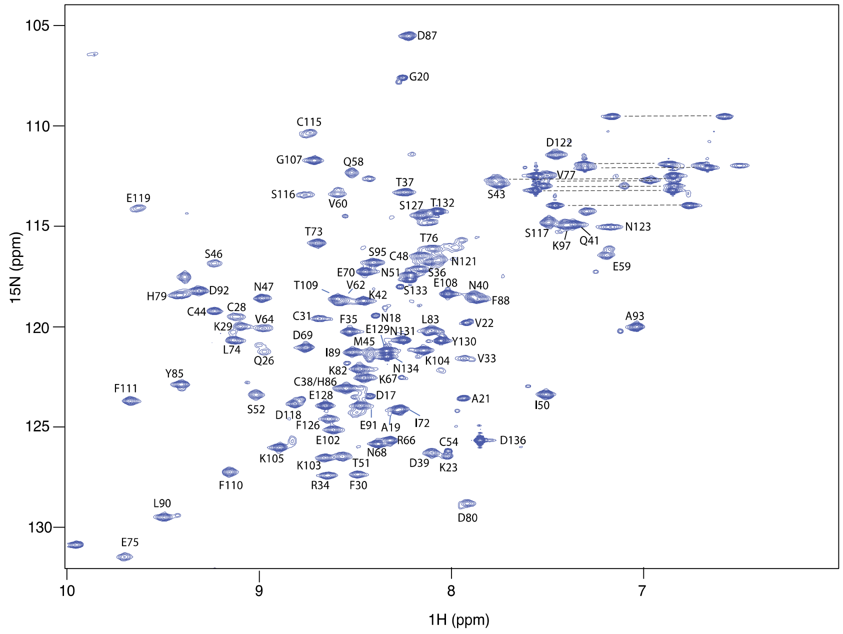

**Figure S10. $^{1}H$-$^{15}N$ HSQC Assignments of TβRI alone and bound to TGM-D2.** A. $^{1}H$-$^{15}N$ HSQC spectra of TβRI alone with peaks assigned. B. $^{1}H$-$^{15}N$ HSQC spectra of TβRI bound to TGM-D2 with peaks assigned. Dashed horizontal lines indicate Nε/Hε pairs. Spectra recorded in 25 mM HEPES, 50 mM sodium chloride, 0.02% azide, 5% $^{2}H_2O$ pH 6.0, 300K

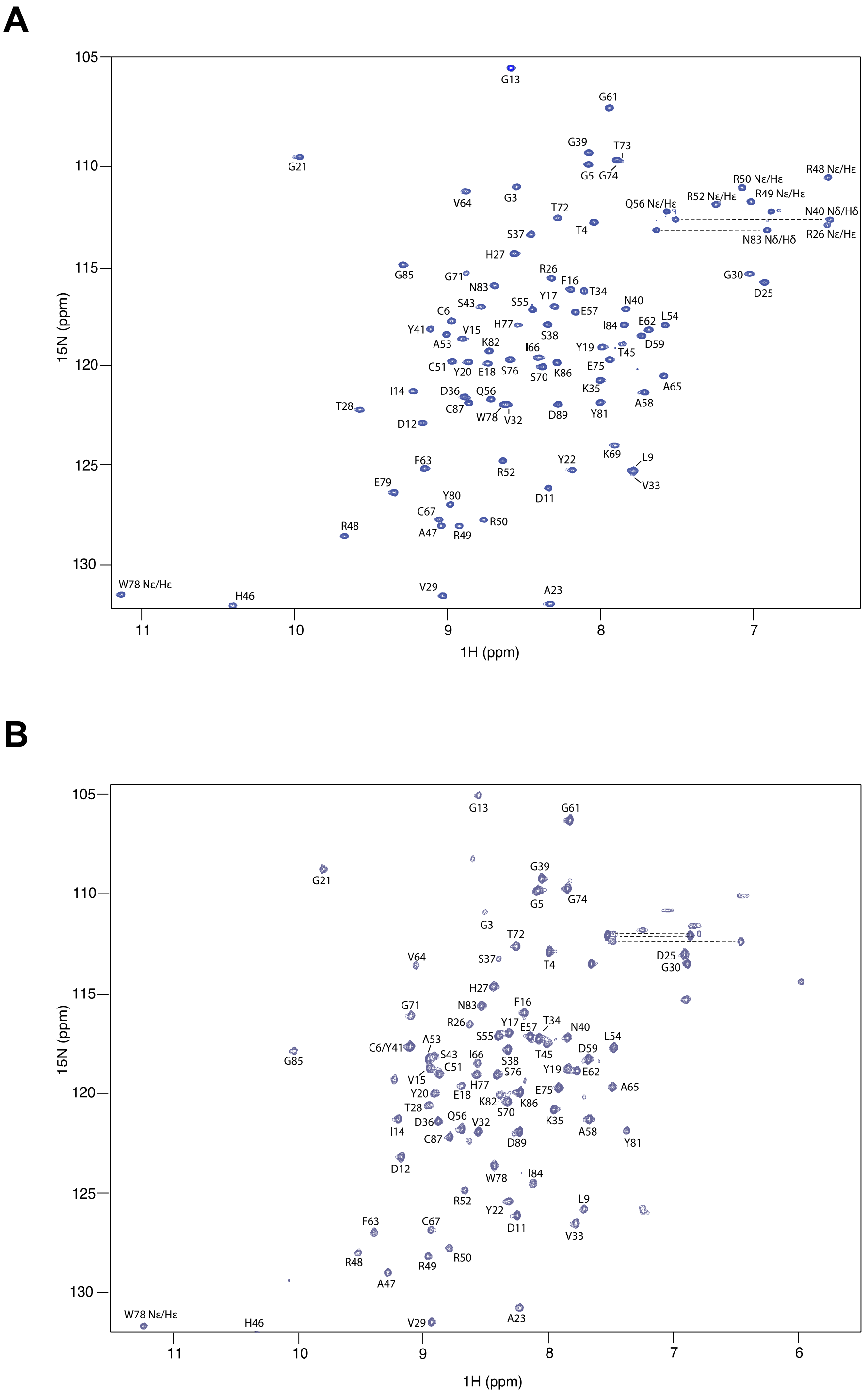

**Figure S11. $^{1}H$-$^{15}N$ HSQC Assignments of TGM-D3 alone and bound to TβRII and predicted secondary structure.** A-B. $^{1}H$-$^{15}N$ HSQC spectra of TGM-D3 alone (A) or bound to TβRII (B) with peaks assigned. Dashed horizontal lines indicate Nε/Hε pairs. C. PECAN-based prediction of TGM-D3 secondary structure. Positive values indicate β–strand probability; negative values indicate helical probability. Spectra recorded in 25 mM sodium phosphate, 50 mM sodium chloride, 5% $^{2}H_2O$ pH 6.0, 310K.

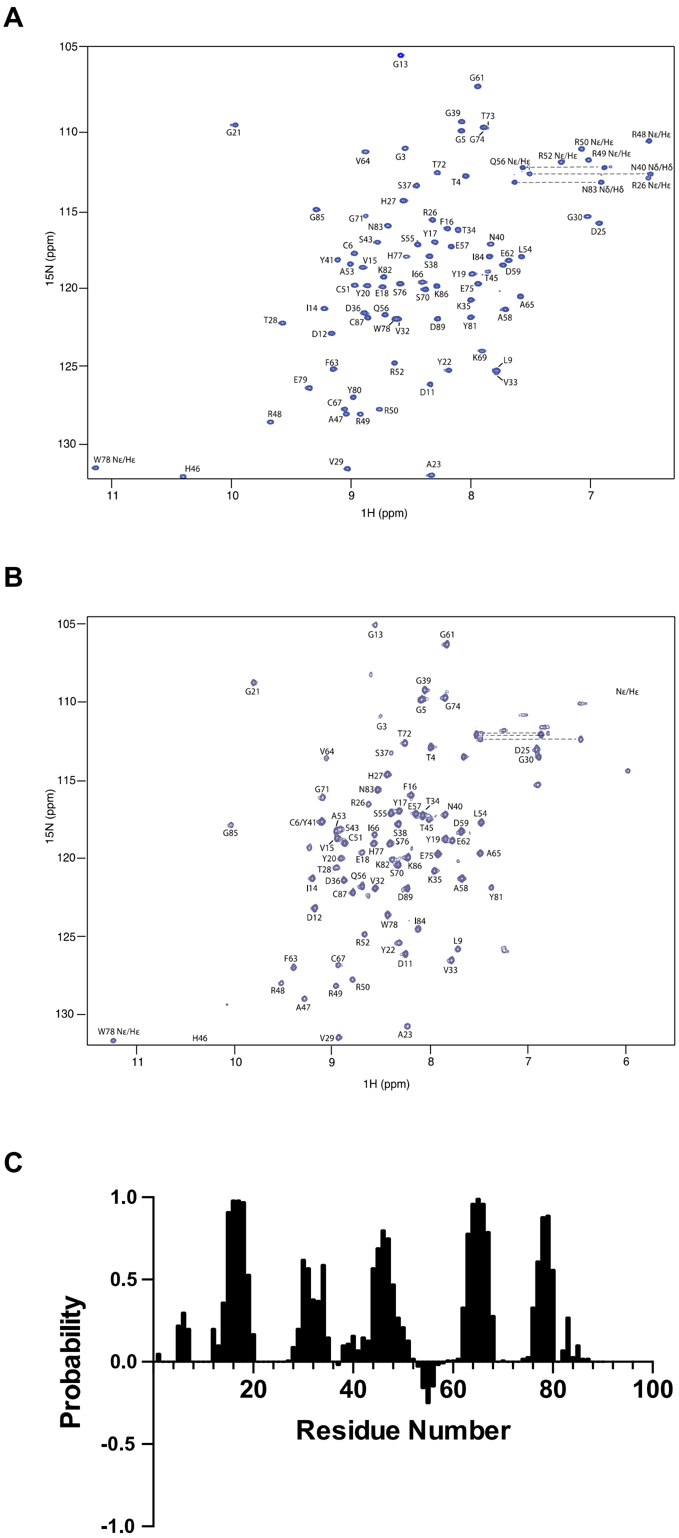

**Figure S12. Alignment of TGM family domains.** A. Alignment of TGM Domains 1-5 with TGM D3. Red indicates conserved residues while blue indicates similar residues. Overlaid on top are the secondary structural features of TGM D3. B. Alignment of TGM Domain 3 with the D3 of TGM-4, -5, and -6. Red indicates conserved residues while blue indicates similar residues. Overlaid on top are the secondary structural features of TGM D3.

A

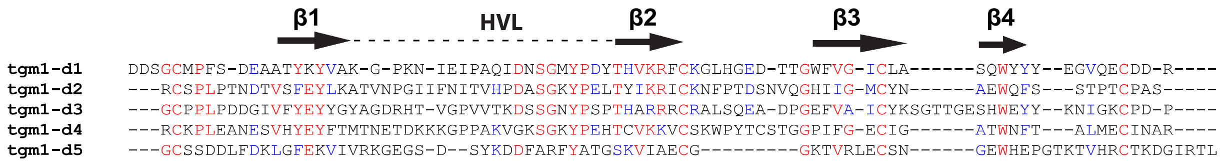


B

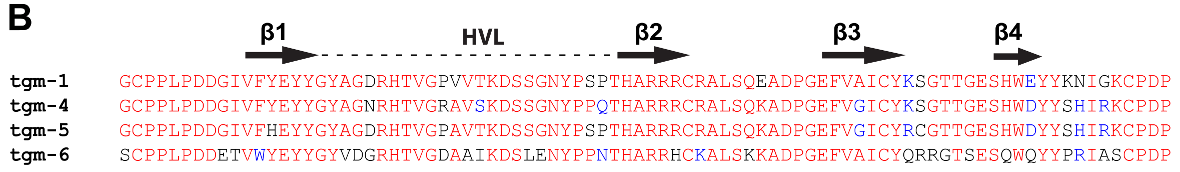

**Figure S13. Secondary Structure Prediction of HpARI and HpBARI.** A Sequence of CCP domains in HpARI. Overlaid on top are the predicted secondary structural features. B. Sequence of CCP domains in HpBARI. Overlaid on top are the predicted secondary structural features.

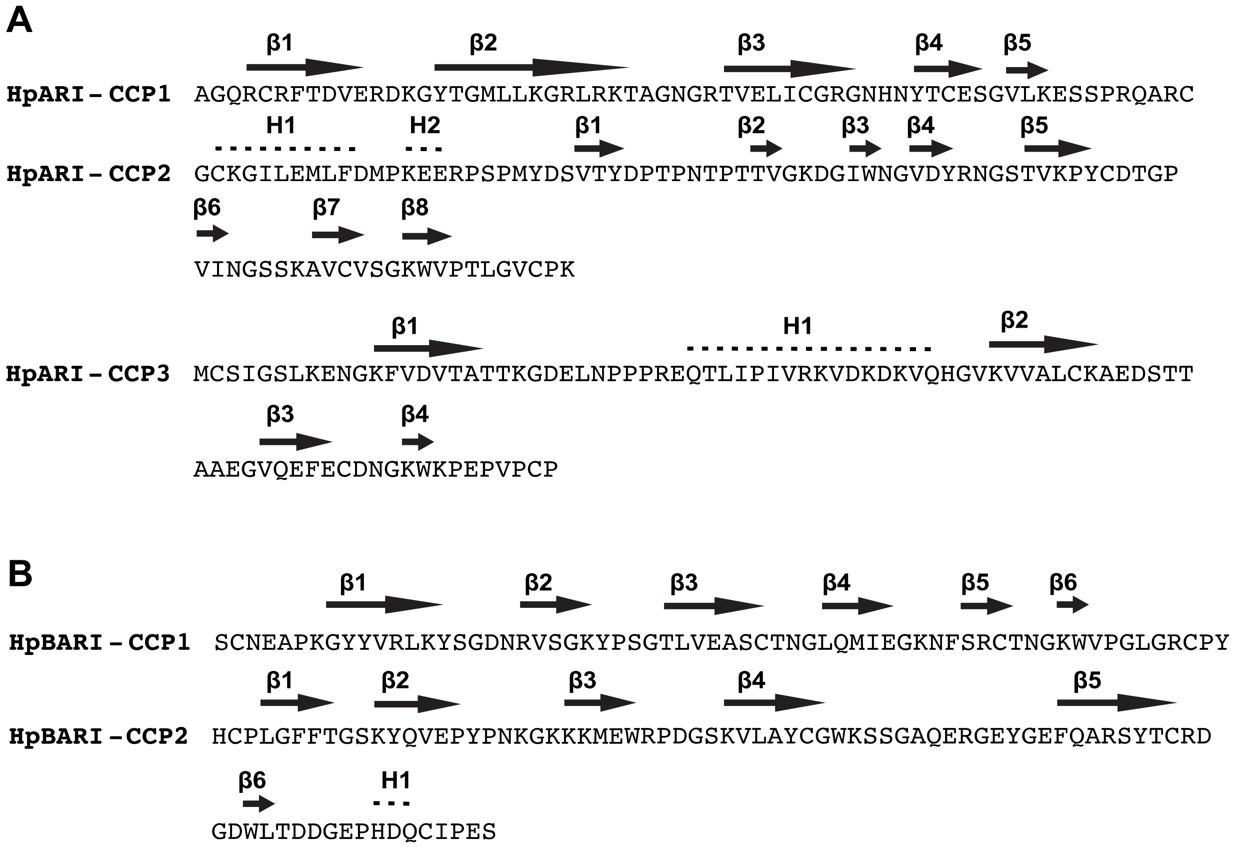
